# Evolutionary fingerprint in rodent PD1 confers weakened activity and enhanced tumor immunity compared to human PD1

**DOI:** 10.1101/2024.09.21.614250

**Authors:** Takeya Masubuchi, Lin Chen, Nimi Marcel, George A. Wen, Christine Caron, Jibin Zhang, Yunlong Zhao, Gerald P. Morris, Xu Chen, Stephen M. Hedrick, Li-Fan Lu, Chuan Wu, Zhengting Zou, Jack D. Bui, Enfu Hui

**Affiliations:** Department of Cell and Developmental Biology, School of Biological Sciences, University of California San Diego, La Jolla, CA 92093; Key Laboratory of Zoological Systematics and Evolution, Institute of Zoology, Chinese Academy of Sciences, Beijing, 100101, China; Department of Molecular Biology, School of Biological Sciences, University of California San Diego, La Jolla, CA 92093; Department of Pathology, University of California San Diego, La Jolla, CA 92093; Department of Neurosciences, University of California, San Diego, La Jolla, CA, 92093; Experimental Immunology Branch, National Cancer Institute, National Institutes of Health, Bethesda, MD 20892, USA

## Abstract

Mechanistic understanding of the immune checkpoint receptor PD1 is largely based on mouse models, but human and mouse PD1 orthologs exhibit only 59.6% identity in amino acid sequences. Here we show that human PD1 is more inhibitory than mouse PD1 due to stronger interactions with the ligands PDL1 and PDL2 and with the effector phosphatase Shp2. A novel motif highly conserved among PD1 orthologs in vertebrates except in rodents is primarily responsible for the differential Shp2 recruitment. Evolutionary analysis suggested that rodent PD1 orthologs uniquely underwent functional relaxation, particularly during the K-Pg boundary. Humanization of the PD1 intracellular domain disrupted the anti-tumor activity of mouse T cells while increasing the magnitude of anti-PD1 response. Together, our study uncovers species-specific features of the PD1 pathway, with implications to PD1 evolution and differential anti-PD(L)1 responses in mouse models and human patients.

## INTRODUCTION

Research conducted on laboratory mice has offered valuable insight into the molecular and cellular mechanisms of immune function. In this regard, mouse models have provided key successes in translational studies, including defining and identifying immune checkpoint inhibitors for use in cancer immunotherapy (*1–6*). Nevertheless, mice and humans differ considerably in the relative abundance of immune cell types, the expression patterns of genes, and the precise functions of certain genes (*7–9*). These differences may explain why checkpoint inhibitor studies in mice have not yet offered substantial insight into the limitations and adverse reactions of immune therapy in human patients(*10–14*). An in-depth understanding of the conservation and divergence of mouse and human immunity at the molecular level is required to better translate findings from mouse models to human trials.

The immune checkpoint receptor PD1 crucially regulates peripheral tolerance and immunity against cancer and infection (*1–5, 15–18*). Although PD1 blocking drugs have demonstrated impressive anti-tumor activity in a subset of human cancer patients (*3, 19–23*), the low response rate and associated immune-related adverse effects, which are often not recapitulated in mice, necessitate a better understanding of PD1 signaling in a human-specific context.

Best known to be expressed by T cells, PD1 contains an immunoglobulin-variable-like (IgV) ectodomain (ECD), a transmembrane domain (TMD), and an intracellular domain (ICD) that harbors two conserved phosphorylatable tyrosines embedded in an immunoreceptor tyrosine inhibitory motif (ITIM) and an immunoreceptor tyrosine switching motif (ITSM), respectively. PDL1, the major ligand of PD1, contains an N-terminal IgV domain that binds to the IgV of PD1 through the front β-face (GFCC’) (*24–26*). This binding triggers PD1 phosphorylation at ITIM and ITSM, which recruits the phosphatase Shp2 to dephosphorylate components in the TCR and costimulatory signaling pathways (*27–30*).

One remarkable, yet largely overlooked feature of PD1 is its relatively low conservation among vertebrates. Whereas intracellular enzymes such as Lck, Zap70, Csk and Shp2 exhibit 95-99% amino acid (AA) sequence identity between human and mouse, human PD1 (*hu*PD1) and mouse PD1 (*mo*PD1) share only 59.6% AA sequence identity, exhibiting substantial differences in their ECD, TMD and ICD. Likewise, many other inhibitory immunoreceptors, including BTLA, Tim3, Lag3, TIGIT, SIRPα, and Siglecs also exhibit low sequence identity between humans and mice, with the exception of CTLA-4, whose ICD is 100% conserved. The cross-species divergence in PD1 sequence suggests that it is still being refined by evolutionary pressure (*31*), but little is known about why, how, and to what extent PD1 function differs across species.

In this study, we quantitatively compared the inhibitory activities of *hu*PD1, *mo*PD1, and their chimeras using biophysical assays, coculture assays and mouse tumor models. These experiments revealed substantial divergence of *hu*PD1 and *mo*PD1 at both biochemical and functional levels. We found that rodent PD1 has a unique pre-ITSM sequence that confers its weaker Shp2 recruitment and less inhibitory function as a consequence of a gradual functional relaxation of rodent PD1 orthologs during evolution. These results demonstrate cross-species differences in the PD1:PDL1 pathway with implications to translation of preclinical findings to human therapeutics.

## RESULTS

### HuPD1 is more inhibitory than moPD1 in both human and mouse T cells

Quantitative comparison of *hu*PD1 and *mo*PD1 activities requires precise control of their expression levels in the same cellular background. We first sought to achieve this in a well-defined coculture system consisting of Jurkat human T cell line (Jurkat) as the responder cell and Raji human B cell line (Raji) as the antigen presenting cell (APC). We established PD1-deficient (*PDCD1*^−/−^) Jurkat cells via CRISPR/Cas9 and transduced them with either *hu*PD1 or *mo*PD1 (**Fig. 1A**), each tagged with a C-terminal monomeric GFP (mGFP), which enabled us to assess the expressions of both PD1 orthologs using a common fluorescent readout, rather than using antibody staining that would require two clones of anti-PD1. Optimization of lentivirus titers allowed us to establish two Jurkat lines that stably expressed similar levels of either *hu*PD1-mGFP or *mo*PD1-mGFP (**Fig. 1A**). The use of the weak dSV40 promoter ensured modest PD1 expression resembling endogenous PD1 expression on human primary CD4^+^ T cells (**fig. S1**). To establish PDL1-expressing APCs, we transduced CD80-deficient (*CD80^−/−^*) Raji cells with either *hu*PDL1-mCherry or *mo*PDL1-mCherry, expressed at comparable amounts indicated by mCherry fluorescence (**Fig. 1A**). CD80 deletion avoided *cis*-CD80:PDL1 interaction, known to block PD1:PDL1 interaction (*32–35*).

**Fig. 1.**
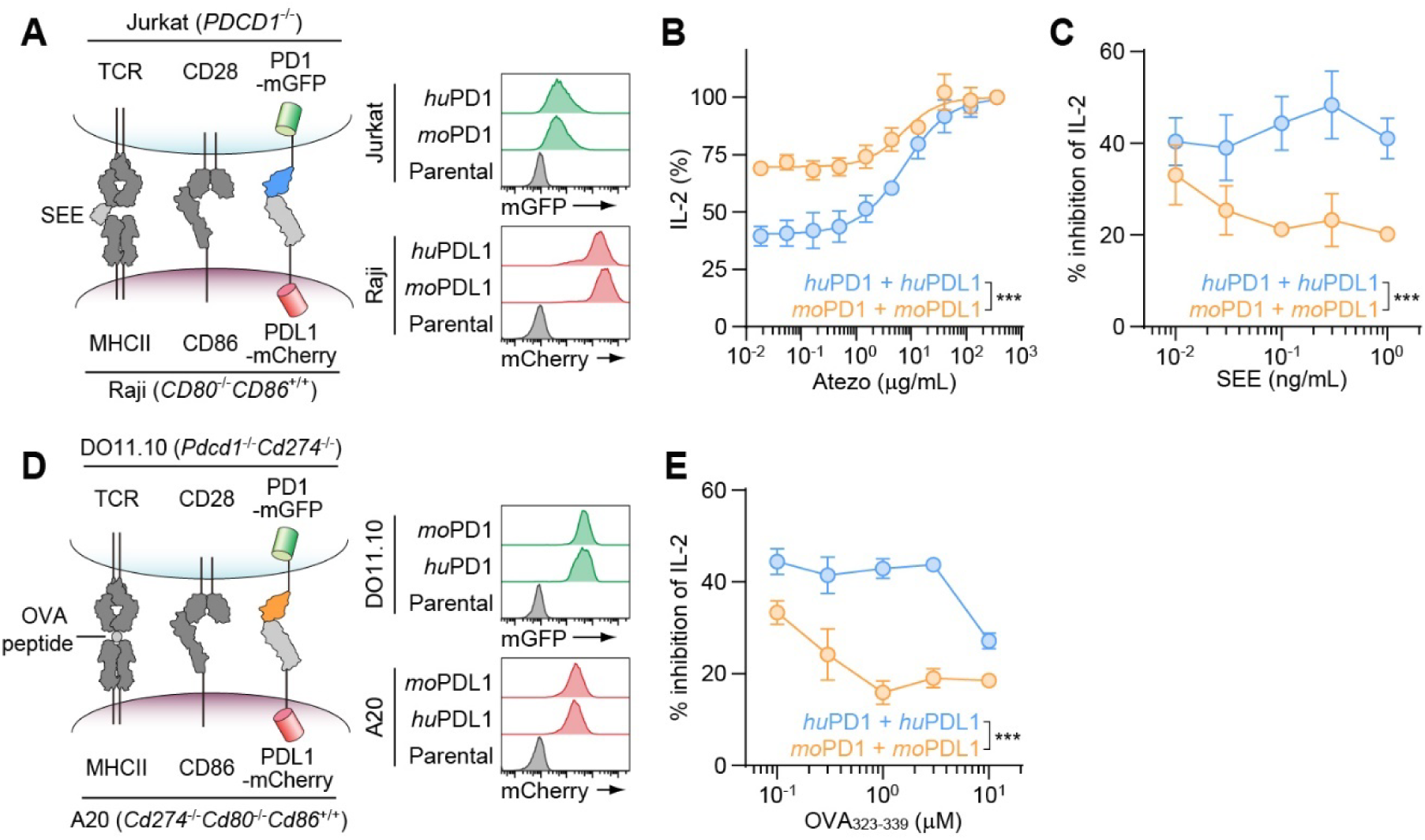
*Hu*PD1:PDL1 interaction more potently inhibited T cell function than *mo*PD1:PDL1 interaction. **(A)** Left, cartoon depicting a human-derived coculture assay containing Jurkat expressing TCR, CD28 and mGFP-tagged *hu*PD1 or *mo*PD1, and SEE-pulsed Raji APCs expressing MHCII, CD86, and mCherry-tagged *hu*PDL1 or *mo*PDL1. Right, FACS histograms showing the expression of indicated PD1-mGFP or PDL1-mCherry variant on the Jurkat or Raji cells. **(B)** Atezolizumab dose-responses of secreted IL-2 in the indicated Jurkat:Raji coculture in the presence of 0.1 ng/mL SEE. Data were normalized to the IL-2 amounts at the highest [atezolizumab]. **(C)** Superantigen dose-responses of % IL-2 inhibition mediated by either *hu*PD1:*hu*PDL1 or *mo*PD1:*mo*PDL1 pair in the Jurkat:Raji coculture assay depicted in **A**. Raji cells were pre-treated with 0 or 120 µg/mL Atezolizumab. **(D)** Left, cartoon depicting a mouse-derived coculture assay containing DO11.10 cells expressing TCR, CD28 and mGFP-tagged *hu*PD1 or *mo*PD1, and OVA_323-339_-pulsed A20 APCs expressing MHCII, CD86, and mCherry-tagged *hu*PDL1 or *mo*PDL1. Right, FACS histograms showing the expression of indicated PD1-mGFP or PDL1-mCherry variant on the DO11.10 or A20 cell. **(E)** OVA_323-339_ dose-responses of % IL-2 inhibition mediated by either *hu*PD1:*hu*PDL1 or *mo*PD1:*mo*PDL1 pair in the DO11.10:A20 coculture assay depicted in **D**. Data are mean ± SD from three independent experiments. ***P < 0.001; two-way ANOVA.

To compare the suppressive activities of *hu*PD1 and *mo*PD1, we set up two coculture assays in parallel with matched species of PD1 and PDL1: (i) Jurkat (*hu*PD1-mGFP) incubated with superantigen (SEE)-loaded Raji (*hu*PDL1-mCherry) to examine *hu*PD1:PDL1 signaling; (ii) Jurkat (*mo*PD1-mGFP) incubated with SEE-loaded Raji (*mo*PDL1-mCherry) to recapitulate *mo*PD1:PDL1 signaling. In both settings, we titrated PD1:PDL1 interactions using atezolizumab, an anti-PDL1 antibody that blocks both *hu*PDL1 and *mo*PDL1 with similar capacities (*36*). Indeed, atezolizumab dose-dependently increased IL-2 production in both settings (**Fig. 1B**). These data revealed that *hu*PD1:PDL1 interaction inhibited IL-2 secretion by ∼60%, while the mouse counterpart did so by only ∼31%. Titration of SEE revealed that *hu*PD1:PDL1 was consistently more suppressive than *mo*PD1:PDL1 across a range of TCR signaling strength (**Fig. 1C**), despite the slightly lower expression of *hu*PDL1 than *mo*PDL1 (**Fig. 1A**). Although Shp2, a known mediator of PD1 signaling, is highly conserved between humans and mice (**fig. S2**), we considered the possibility that the human-derived Jurkat environment might be less permissive to *mo*PD1, thereby mitigating its inhibitory activity. We therefore performed a complementary experiment using DO11.10 mouse T cell hybridoma (DO11.10 cells) cocultured with A20 mouse B cell lines (A20 cells), which can present ovalbumin (OVA)-derived peptide 323-339 (OVA_323-339_) via class-II major histocompatibility complex (MHC-II) I-A^d^ (**Fig. 1D**). We transduced *Pdcd1*^−/−^*Cd274*^−/−^ DO11.10 cells with *hu*PD1-mGFP or *mo*PD1-mGFP, and *Cd274*^−/−^*Cd80*^−/−^*Cd86*^+/+^ A20 cells with *hu*PDL1-mCherry or *mo*PDL1-mCherry. Using fluorescence activated cell sorting (FACS), we obtained two lines of DO11.10 cells expressing similar amounts of *hu*PD1-mGFP and *mo*PD1-mGFP, and two lines of A20 cells expressing comparable amounts of *hu*PDL1-mCherry and *mo*PDL1-mCherry (**Fig. 1D**, **fig. S3A, B**). Coculture assays showed that even in the mouse-derived DO11.10 and A20 cells, the *hu*PD1:PDL1 interaction was more suppressive than *mo*PD1:PDL1 interaction (**Fig. 1E**). Thus, when stimulated by comparable amounts of cognate PDL1, *hu*PD1 is more inhibitory than *mo*PD1 in T cells of both human and mouse origin.

### Both ECD and ICD contribute to the stronger inhibitory function of huPD1

*Hu*PD1 and *mo*PD1 share 61.2%, 33.3%, and 58.0% AA identities in ECD, TMD, and ICD, respectively. We next asked which domain(s) of PD1 underlie(s) the different inhibitory activities of *hu*PD1 and *mo*PD1 using PD1 chimeras. We created Jurkat cell lines expressing similar levels of *hu*PD1, *mo*PD1, or chimeric PD1 in which the ECD, TMD or ICD of *hu*PD1 was swapped with the corresponding domain of *mo*PD1, designated as *hu*PD1*^mo^*^ECD^, *hu*PD1*^mo^*^TMD^ and *hu*PD1*^mo^*^ICD^, respectively (**Fig. 2A**). We then cocultured each Jurkat cell line with SEE-pulsed *CD80*^−/−^*CD86*^+/+^ Raji expressing either *hu*PDL1-mCherry or *mo*PDL1-mCherry to ensure matched species of PDL1 and the ECD species of PD1 chimera. These experiments showed that swapping either the ECD or the ICD, but not the TMD decreased the abilities of *hu*PD1 to inhibit both IL-2 secretion and CD69 expression (**Fig. 2B**). These results suggest that both the ECD and the ICD contribute to the more suppressive activity of *hu*PD1.

**Fig. 2.**
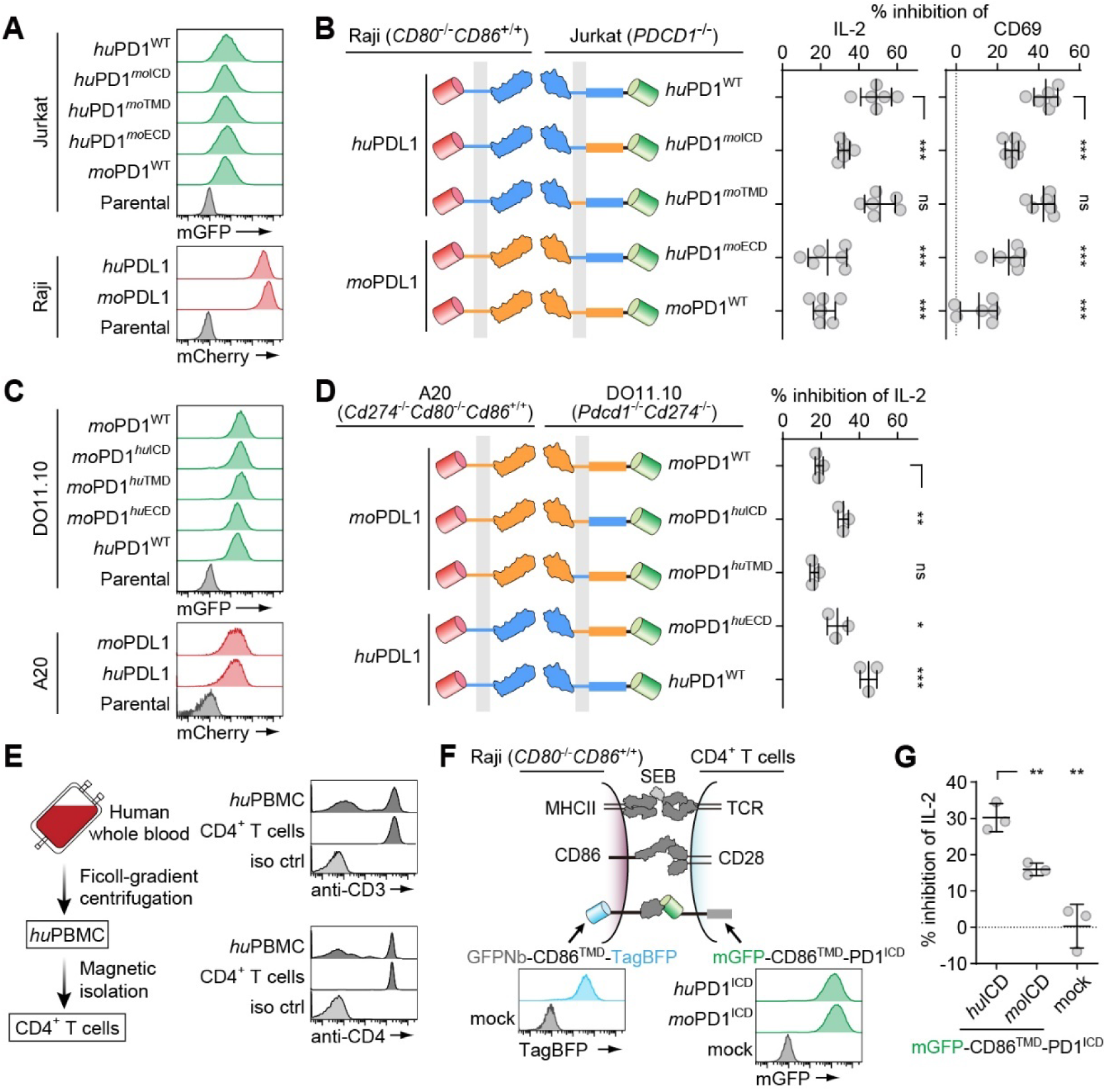
Both ECD and ICD of *hu*PD1 contribute to its stronger inhibitory function than that of *mo*PD1. **(A)** FACS histograms showing the expressions of indicated mGFP-tagged PD1 or mCherry-tagged PDL1 variants on Jurkat or Raji cells used for cocultures. **(B)** Left, a cartoon depicting the PD1:PDL1 pairs tested in five parallel Jurkat:Raji coculture assays. Right, bar graphs showing the % inhibition of IL-2 secretion and of CD69 expression by the PD1:PDL1 pairs indicated on the left (n = 6 independent experiments). **(C)** FACS histograms showing the expressions of indicated mGFP-tagged PD1 or mCherry-tagged PDL1 variants on DO11.10 or A20 cells used for cocultures. **(D)** Left, a cartoon depicting the PD1:PDL1 pairs tested in five parallel A20:DO11.10 coculture assays. Right, a bar graph showing the % inhibition of IL-2 secretion by the PD1:PDL1 pairs indicated on the left (n = 3 independent experiments). **(E)** Left, diagram showing the purification method for human primary CD4^+^ T cells. Right, FACS histograms of CD3 and CD4 expressions on *hu*PBMC and CD4^+^ T cells. (F) Top, cartoon depicting a coculture assay containing CD4^+^ T cells expressing TCR, CD28, and mGFP^ECD^-TMD-PD1^ICD^, and SEB-pulsed Raji APCs expressing MHCII, CD86, and GFPNb-TM-TagBFP. Bottom, FACS histograms showing the expression of indicated PD1 chimera or GFPNb-TM-TagBFP on CD4^+^ T or Raji cells, respectively. (G) A dot plot showing % inhibition of IL-2 secretion by the PD1 chimera:GFPNb association in CD4^+^ T cell:Raji coculture depicted in **F** (n = 3 independent experiments). Data are mean ± SD. *P < 0.05; **P < 0.01; ***P < 0.001; ns, not significant; student’s t-test.

In a reciprocal set of experiments, we compared the inhibitory activities of PD1 chimeras in the DO11.10:A20 mouse cell culture system, but used *mo*PD1 as a template to construct three chimeras: *mo*PD1*^hu^*^ECD^, *mo*PD1*^hu^*^TMD^ and *mo*PD1*^hu^*^ICD^, with either the ECD, TMD or ICD of *mo*PD1 replaced by the corresponding domain of *hu*PD1. We generated five DO11.10 cell lines expressing comparable amounts of mGFP-tagged *hu*PD1, *mo*PD1 or each of the three foregoing PD1 chimeras (**Fig. 2C**), and stimulated each with peptide-pulsed A20 cells expressing PDL1 whose species matched the ECD of the corresponding PD1 variant. Consistent with the results in Jurkat:Raji system, both ECD and ICD contributed to the stronger activity of the human orthologs (**Fig. 2D**).

To validate the differential inhibitory capacities of *hu*PD1 and *mo*PD1 in primary T cells, we transduced human CD4^+^ T cells with either *hu*PD1 or *mo*PD1, each with the ECD replaced by mGFP, and expressed at similar levels (**Fig. 2, E** and **F**). To trigger GFP-PD1 chimeras without stimulating endogenous PD1, we created *CD80*^−/−^*CD86*^+/+^ Raji APCs expressing a GFPNb-TM-tagBFP, in which a GFP nanobody was fused to CD86TMD and a C-terminal tagBFP. Upon coculturing GFP^+^CD4^+^ T cells with SEB-pulsed *CD80^−/−^CD86^+/+^GFPNb-TM-tagBFP*^+^ Raji APCs (**Fig. 2F**), GFP^ECD^-*hu*PD1^ICD^ and GFP^ECD^-*mo*PD1^ICD^ inhibited IL-2 production by 30% and 16%, respectively, compared to conditions using *CD80^−/−^CD86^+/+^*Raji APCs lacking GFPNb-TM-tagBFP (**Fig. 2G**). This result further supported the notion that *hu*PD1 ICD is intrinsically more suppressive than *mo*PD1 ICD.

### HuPD1:PDL1/2 interaction is stronger than moPD1:PDL1/2 interaction

Having shown that the stronger inhibitory activity of *hu*PD1 mapped in part to the ECD (**Fig. 2**, *hu*PD1^WT^ versus *hu*PD1*^mo^*^ECD^, *mo*PD1*^hu^*^ECD^ versus *mo*PD1^WT^), we next compared the 3D affinities of *hu*PD1 and *mo*PD1 with their respective cognate ligands using bio-layer interferometry (BLI). *hu*PD1^ECD^ or *mo*PD1^ECD^ dose-dependently bound sensor-immobilized, species-matched PDL1^ECD^ or PDL2^ECD^ (**Fig. 3A**). The resulting dissociation constants (*K*_d_) showed a 3.2-fold higher affinity for the human PD1:PDL1 pair than for the mouse counterpart, and a 24-fold higher affinity for the human PD1:PDL2 pair than for the mouse counterpart (**Fig. 3B**). These results are in qualitative agreement with surface plasmon resonance measurements using refolded bacterially expressed proteins (*37*). Moreover, in a cell-free membrane reconstitution assay (*38, 39*), we found that *hu*PD1:*hu*PDL1 interactions drove more association of large unilamellar vesicles (LUVs) with supported lipid bilayers (SLBs) than did *mo*PDL1:*mo*PD1 interactions (**fig. S4**), indicating a higher 2D affinity for the human pair.

**Fig. 3.**
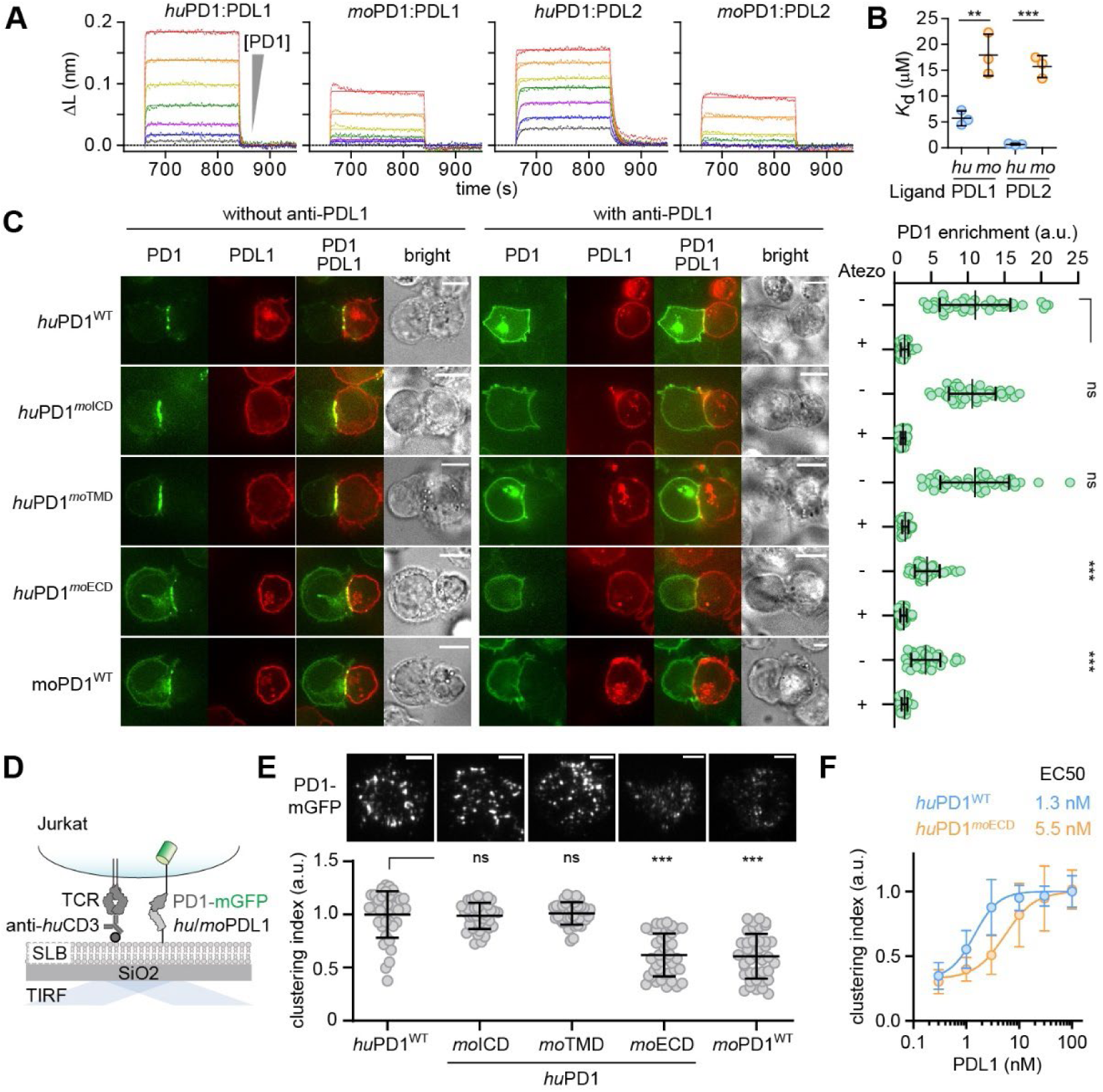
Human PD1 ECD associates with its ligands more strongly than mouse PD1 ECD. **(A)** Representative BLI data obtained for the indicated PD1:ligand pairs with the PDL1 or PDL2 immobilized on the sensor chip and increasing concentrations of PD1-ECD presented in the solution. **(B)** *K*_d_ values of the indicated PD1:ligand pairs based on the three independent BLI experiments. **(C)** Left, representative confocal images of a conjugate between Jurkat expressing the indicated PD1 variants (green) and Raji expressing the indicated PDL1 variants (red) with or without anti-PDL1 (atezolizumab). Right, a dot plot showing the synaptic enrichment score of each PD1 variant in the presence or absence of atezolizumab (n = 40 cells). **(D)** A cartoon depicting a cell-SLB assay, in which Jurkat cells expressing mGFP-tagged PD1 variant interacted with a SLB functionalized with anti-*hu*CD3ε and *hu*/*mo*PDL1-His. PD1 microclusters were visualized via TIRF-M. **(E)** Top, representative TIRF images showing microclusters of indicated mGFP-tagged PD1 variant in a Jurkat cell that contacted the SLB functionalized with 3 nM *hu*PDL1 or *mo*PDL1 that matched the species of the ECD of the corresponding PD1 variant. Bottom, dot plots showing the clustering indices of all five PD1 variants (See methods) (n = 40 cells). (F) Dose-response curves showing PD1 clustering indices of the indicated PD1 variant plotted against [PDL1] (n = 3 independent experiments). Scale bars: 5 µm. Data are mean ± SD. **P < 0.01; ***P < 0.001; ns, not significant; student’s t-test.

We further compared human and mouse PD1:PDL1 interactions using full length *hu*PD1, *mo*PD1 and their chimeras. To examine PD1:PDL1 *trans*-interaction (*38, 39*), we measured PDL1-dependent enrichment of PD1 to the APC:T-cell interface in a Jurkat:Raji coculture assay. After incubation of Jurkat cells expressing mGFP-tagged *hu*PD1, *mo*PD1 or their chimeras with SEE-loaded Raji APCs (*CD80*^−/−^*CD86*^+/+^) expressing either *hu*PDL1-mCherry or *mo*PDL1-mCherry, we observed more accumulation of PD1 variants with the human ECD (*hu*PD1^WT^, *hu*PD1*^mo^*^TMD^, *hu*PD1*^mo^*^ICD^) to the Jurkat:Raji border compared to PD1 variants containing the mouse ECD (*hu*PD1*^mo^*^ECD^ and *mo*PD1^WT^) (**Fig. 3C**). As expected, the presence of atezolizumab abrogated the synaptic enrichment of PD1 (**Fig. 3C**). We extended these findings in a PD1 microcluster assay (*27*) that employs SLBs as artificial APCs (**Fig. 3D**). When the aforementioned Jurkat cells landed onto SLBs containing Okt3 (anti-*hu*CD3ε), human ICAM1 (*hu*ICAM1), and equal amounts of *hu*PDL1 or *mo*PDL1, PD1 variants containing *hu*PD1 ECD (*hu*PD1^WT^, *hu*PD1*^mo^*^TMD^, and *hu*PD1*^mo^*^ICD^) formed more intense microclusters than PD1 variants containing mouse ECD (*hu*PD1*^mo^*^ECD^ and *mo*PD1^WT^), recorded by total internal reflection fluorescence microscopy (TIRF-M) (**Fig. 3E**, **fig. S5** and **supplementary movies 1** and **2**). Titration of PDL1 on SLB revealed that *hu*PD1*^mo^*^ECD^ required 4.2-fold more cognate ligand to cluster to a similar degree as did *hu*PD1^WT^ (**Fig. 3F**).

Collectively, data in this section demonstrate that *hu*PD1 binds cognate PDL1 and PDL2 more strongly than does *mo*PD1.

### HuPD1 recruits Shp2 more strongly than moPD1

We then turned our attention to the ICD, since our chimera experiments showed that *hu*PD1 ICD mediates stronger inhibitory effects than *mo*PD1 ICD (**Fig. 2**, *hu*PD1^WT^ versus *hu*PD1*^mo^*^ICD^, *mo*PD1*^hu^*^ICD^ versus *mo*PD1^WT^). The ICD of PD1 is known to recruit Shp2 (*27, 40*). Indeed, treatment of T cell:APC co-cultures with SHP099, an allosteric inhibitor of Shp2 (*41*), decreased the ability of PD1 to inhibit IL-2 production by about 50%, in both human and mouse CD8+ T cells (**fig. S6**). To determine the biochemical mechanism underlying the differential suppressive activities of *hu*PD1 ICD and *mo*PD1 ICD, we compared their abilities to recruit Shp2 by imaging PD1:Shp2 association in T cells in a cell-SLB system (**Fig. 4A**). After incubating Jurkat cells expressing *hu*PD1^WT^-mGFP or *hu*PD1*^mo^*^ICD^-mGFP with *hu*PDL1-functionalized SLBs, we fixed and permeabilized the SLB-associated cells, and stained endogenous Shp2 with AlexaFluor-647 anti-Shp2 (anti-Shp2*AF647) (**Fig. 4A**). TIRF-M detected intense microclusters for both *hu*PD1^WT^-mGFP and *hu*PD1*^mo^*^ICD^-mGFP, as they both contained the ECD of *hu*PD1 However, *hu*PD1^WT^-mGFP recruited significantly more Shp2 than did *hu*PD1*^mo^*^ICD^-mGFP (**Fig. 4B**). Likewise, when human primary CD4^+^ T cells transduced with mGFP-tagged *hu*PD1^WT^, *hu*PD1*^mo^*^ICD^, or tailless *hu*PD1 (*hu*PD1^TL^) contacted SLBs containing anti-*hu*CD3ε, *hu*PDL1, and *hu*ICAM1, *hu*PD1^WT^ microclusters recruited more Shp2 than did *hu*PD1*^mo^*^ICD^ in CD4^+^ T cells, either based on endogenous Shp2 immunostaining or transduced mCherry-Shp2 (**Fig. 4C**, **fig. S7**). The dim anti-Shp2 signal detected within *hu*PD1^TL^ foci was likely due to endogenous PD1. We further compared the abilities of *hu*PD1 ICD and *mo*PD1 ICD to recruit mouse Shp2 in mouse cells. When DO11.10 cells expressing *mo*PD1^WT^-mGFP or *mo*PD1*^hu^*^ICD^-mGFP contacted SLBs harboring anti-*mo*CD3ε, *mo*PDL1, *mo*ICAM1 (**Fig. 4D**), both PD1 variants formed microclusters, but *mo*PD1*^hu^*^ICD^ recruited more Shp2 than did *mo*PD1^WT^, as indicated by anti-Shp2*AF647 staining (**Fig. 4E**). Notably, Shp2 is highly conserved between human and mouse, especially the PD1 binding module tandem SH2 domains (tSH2), exhibiting 100% sequence identity between the two species (**fig. S2**). Thus, *hu*PD1 is more capable of recruiting the intracellular effector Shp2 than *mo*PD1.

**Fig. 4.**
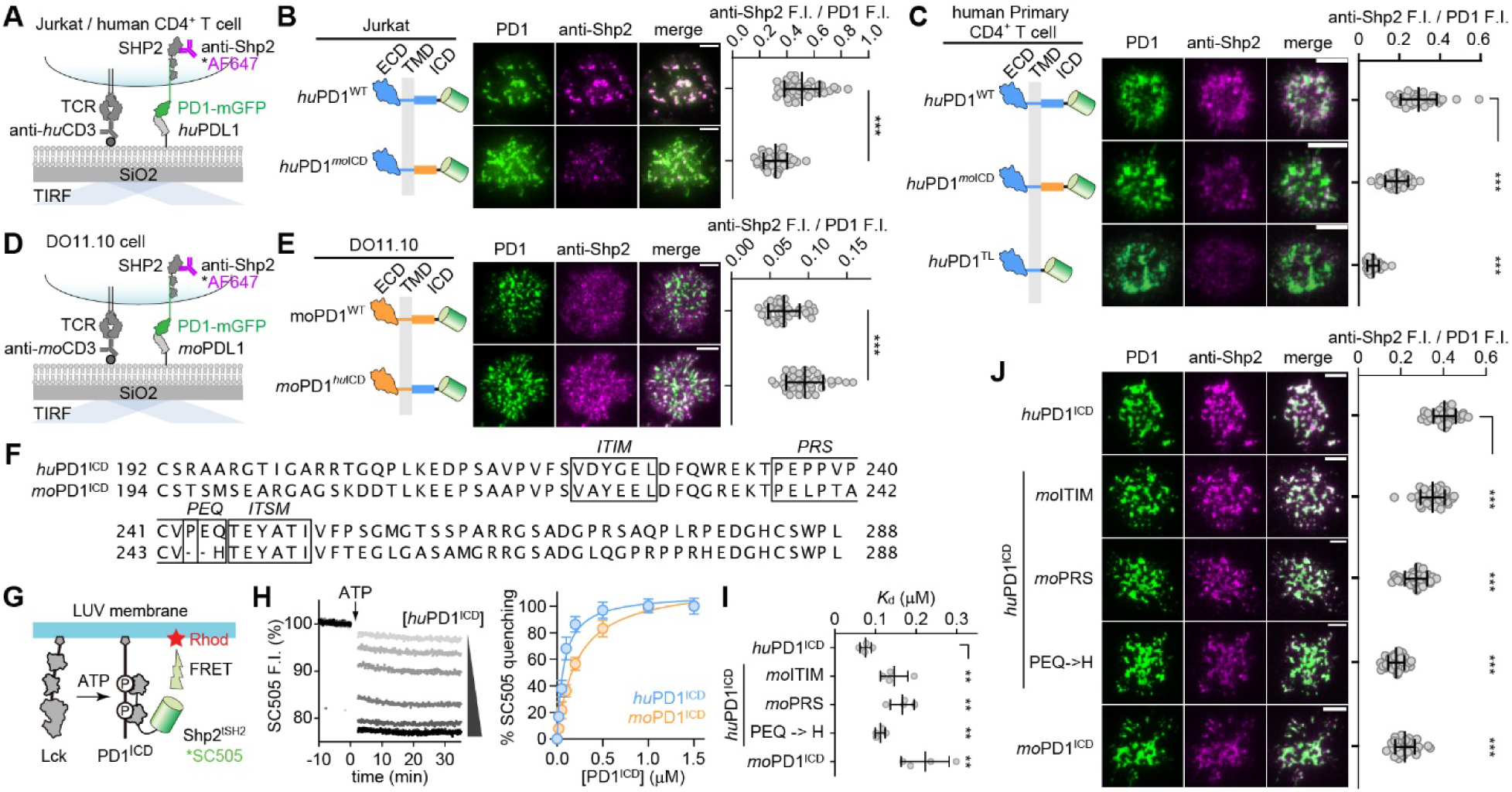
*Hu*PD1 more strongly recruits Shp2 than does *mo*PD1 due to three non-conserved motifs. (**A**) A cartoon showing a cell-SLB assay for TIRF imaging of PD1:Shp2 association in Jurkat or human CD4^+^ T cells. (**B**) Left, representative TIRF images of the indicated mGFP-tagged PD1 variants (green) and anti-Shp2 (magenta) at the interface of a Jurkat cell and the SLB as depicted in **A**. Right, dot plots showing the anti-Shp2 F.I. normalized to PD1 F.I. (n = 40 cells). (**C**) Same as **B**, except that human CD4^+^ T cells expressing the indicated PD1 variants were imaged (n = 40 cells). (**D, E**) Same as **A, B**, except that DO11.10 cells were observed. (**F**) AA sequence alignment of *hu*PD1^ICD^ and *mo*PD1^ICD^, with ITIM, PRS, PEQ/H, and ITSM highlighted. (**G**) Cartoon of a liposome reconstitution assay for measuring PD1^ICD^:Shp2^tSH2^ interaction. (**H**) Left, representative time courses of SC505 (Shp2^tSH2^) F.I. at increasing concentrations of *hu*PD1^ICD^. Right, % SC505 quenching 30 min after ATP addition plotted against the [*hu*PD1^ICD^] and [*mo*PD1^ICD^] (n = 3 independent experiments). (**I**) Bar graphs summarizing apparent *K*_d_ of Shp2^tSH2^ interaction with indicated PD1^ICD^ variants determined via assays shown in **G** and **H** (n = 3 independent experiments). (**J**) Left, representative TIRF images showing Shp2 recruitment to microclusters of indicated PD1 variants in a Jurkat-SLB assays. Right, dot plots showing anti-Shp2 F.I. normalized to PD1 F.I. (n = 40 cells). Scale bars: 5 µm. Data are mean ± SD. **P < 0.01; ***P < 0.001; ns, not significant; student’s t-test.

### A pre-ITSM PEQ motif in huPD1 drives stronger Shp2 recruitment

The stronger Shp2 recruitment by *hu*PD1 than by *mo*PD1 was unexpected because ITSM, known as the major Shp2 docking site, is identical in *hu*PD1 and *mo*PD1. We next sought to identify the molecular determinants in PD1 ICD underlying the differential Shp2 recruitment activities of human and mouse orthologs. Despite the conserved ITSM, PD1 shows variations in the pre-ITSM regions: *hu*PD1 contains an identifiable proline rich sequence (PRS) and a Pro-Glu-Gln (PEQ) sequence immediately upstream to ITSM; *mo*PD1 differs from *hu*PD1 by two residues in the ITIM, four in the PRS, and replaces the PEQ motif with a single His (**Fig. 4F**). To determine whether these motifs contribute to Shp2 recruitment, we constructed, expressed and purified His-tagged *hu*PD1 ICD WT and mutants with each of the three motifs (ITIM, PRS, PEQ) replaced with the corresponding segment in *mo*PD1, designated as *hu*PD1-ICD*^mo^*^ITIM^, *hu*PD1-ICD*^mo^*^PRS^, *hu*PD1-ICD^PEQ->H^, respectively. We then measured the abilities of these proteins to recruit Shp2^tSH2^ using a LUV reconstitution assay with a Förster Resonance Energy Transfer (FRET) readout, as described (**Fig. 4G**) (*42*). We coupled each of the five His-PD1 ICD variants together with His-tagged Lck kinase to rhodamine (energy acceptor)-labeled LUVs, and presented SNAP-cell-505 (SC505, energy donor)-labeled Shp2^tSH2^ in the solution. Subsequent ATP addition triggered PD1 ICD phosphorylation by Lck and recruitment of Shp2^tSH2^*SC505, leading to FRET-mediated SC505 quenching by rhodamine (**Fig. 4G**) in a PD1 ICD dose-dependent fashion, allowing us to calculate the apparent *K*_d_ values (**Fig. 4H**). These experiments revealed a 2.9-fold stronger Shp2 affinity for *hu*PD1-ICD^WT^ than for *mo*PD1-ICD^WT^ (76 nM *K*_d_ for *hu*PD1 and 220 nM *K*_d_ for *mo*PD1), consistent with data in cells (**Figs. 4, B, C** and **E**). Likewise, Shp1, reported to mediate PD1 signaling upon Shp2 deletion (*43*), also bound to *hu*PD1-ICD^WT^ with a higher affinity than to *mo*PD1-ICD^WT^ (**fig. S8**). Swapping ITIM, PRS, or PEQ of *hu*PD1 with the corresponding motif in *mo*PD1 decreased the Shp2 affinity, as reflected by greater *K*_d_ values for *hu*PD1-ICD*^mo^*^ITIM^, *hu*PD1-ICD*^mo^*^PRS^, and *hu*PD1-ICD^PEQ->H^ than for *hu*PD1-ICD^WT^ (**Fig. 4I**). These results suggest that besides the conserved ITSM, *hu*PD1:Shp2 interaction is contributed by three non-conserved motifs – ITIM, PRS, and PEQ – that are either weakened or lacking in *mo*PD1.

To validate the results of LUV-FRET assays in a cellular context, we transduced Jurkat cells with mGFP-tagged *hu*PD1 variants in which ITIM, PRS, or PEQ motif was replaced with the corresponding motif in *mo*PD1, and measured Shp2 recruitment to microclusters of these PD1 mutants (**Fig. 4A**). Consistent with the LUV-FRET assay, all three domain-swapping mutants exhibited weaker Shp2 recruitment compared to *hu*PD1^WT^ (**Fig. 4J**). The relative magnitude of the effect differed in the LUV-FRET assay and the cellular assay, likely due to the use of a truncated form (tSH2) of Shp2, or the lack of other cellular factors in the LUV-FRET assay. In the cellular assay, PEQ->H mutation produced the largest decrease in Shp2 recruitment, to a similar extent as did *hu*PD1*^mo^*^ICD^, which had the entire ICD replaced by the mouse version (**Fig. 4J**). Altogether, data in this section demonstrated that despite the identical ITSM, *hu*PD1 is superior to *mo*PD1 in Shp2 recruitment, largely attributable to the PEQ motif that is substituted by a His in *mo*PD1.

### Rodent PD1 orthologs share a weak pre-ITSM motif

The ability of PEQ->H mutation to markedly reduce Shp2 recruitment motivated us to investigate how this motif diverges among vertebrates. We created a phylogenetic tree based on the AA sequence alignment of full-length PD1 from 236 vertebrates, including amphibians, reptiles, birds, and mammals, color coded based on the pre-ITSM sequence (**Fig. 5A**, see **Methods**). The PEQ motif found in *hu*PD1 is conserved in 131 species. PD1 orthologs in an additional 71 species contain PEQ-like motifs at the corresponding region (**Fig. 5A**). In contrast, PD1 orthologs in the entire rodent clade uniquely contain a lone His at the pre-ITSM position (**Fig. 5A**).

**Fig. 5.**
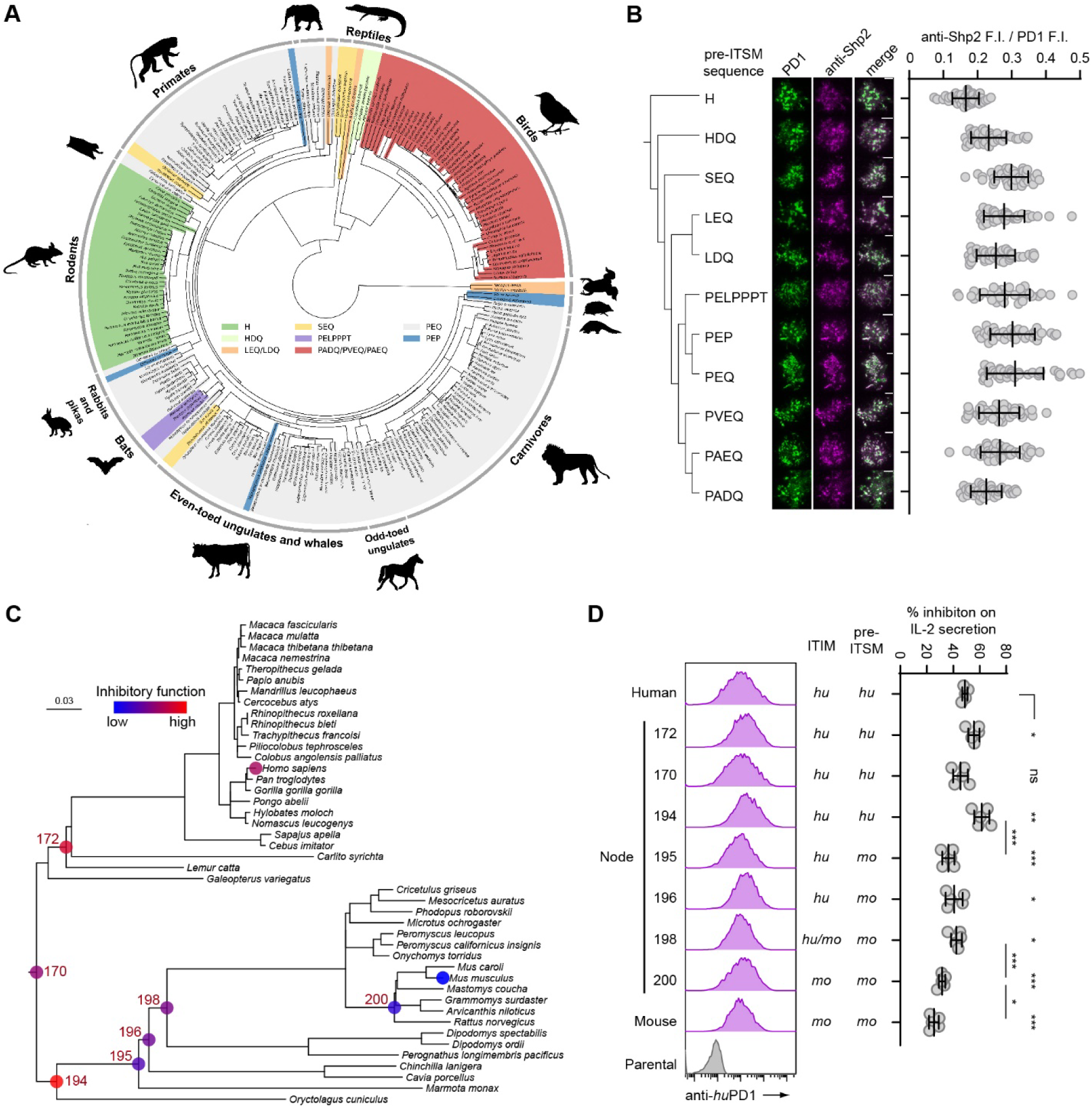
Rodent PD1 orthologs share a unique pre-ITSM sequence that weakens their ability to recruit Shp2. (**A**) Phylogeny of 236 vertebrate species, color-coded based on the pre-ITSM sequence of PD1. (**B**) Left, a phylogenetic tree of pre-ITSM sequence found in PD1 orthologs. Middle, representative TIRF images of microclusters of GFP-tagged *hu*PD1 variant bearing the indicated pre-ITSM sequence (green) and endogenous Shp2 (magenta) in an anti-*hu*CD3ε/*hu*PDL1 stimulated Jurkat cell. Right, dot plots showing anti-Shp2 F.I. normalized to PD1 F.I. in Jurkat cells expressing the indicated *hu*PD1 variant (n = 40 cells). (**C**) The subtree of the 107-species mammal phylogeny spanning the rodent clade and the primate clade. The branch lengths were inferred based on PD1 codon sequence evolution. The marked nodes are mouse, human or the ones with reconstructed ancestral sequences. The color codes show the inhibition ability of the sequences at respective nodes, based on data in D. (**D**) Left, FACS histograms showing the expressions of *hu*PD1 chimera harboring an ICD corresponding to the indicated node, on Jurkat cells. Right, % inhibition of IL-2 secretion by the indicated *hu*PD1 chimera upon stimulating Jurkat (*hu*PD1 chimera) with Raji (*hu*PDL1). The ITIM and pre-ITSM type is labeled: *hu*, human-like, *mo*, mouse like (n = 5 independent experiments). Data are mean ± SD. Scale bars: 5 µm. *P < 0.05; **P < 0.01; ***P < 0.001; ns, not significant; student’s t-test.

We next determined the biochemical consequences of different pre-ITSM sequences in a cell-SLB set up (**Fig. 4A**). We transduced Jurkat cells with *hu*PD1 mutants in which the PEQ motif was replaced with the pre-ITSM motif found in other PD1 orthologs, and measured Shp2 recruitment to PD1 microclusters in anti-*hu*CD3ε/*hu*ICAM1/*hu*PDL1 stimulated Jurkat cells (**Fig. 5B**). TIRF-M showed that *hu*PD1 with rodent pre-ITSM sequence (His) recruited significantly less Shp2 than did *hu*PD1 with PEQ or PEQ-like pre-ITSM sequences.

We further determined to what extent pre-ITSM sequences correlate with the strength of PD1 signaling across five mammalian species: three non-rodent species (human, monkey, dog) that contain the PEQ motif, and two rodents (mouse, rat) that contain an His (**fig. S9A**). To directly compare intracellular signaling of the foregoing PD1 orthologs, we created four PD1 chimeras in which the ICD of *hu*PD1 was replaced with the ICD of monkey, dog, mouse or rat PD1 orthologs. We then engineered five Jurkat cell lines expressing similar levels of either *hu*PD1 or each of the four chimeras. Notably, these PD1 variants differed only in the species of their ICDs, but shared the same ECD and TMD (human), and therefore could be triggered by a single type of Raji cells that expressed *hu*PDL1, and blocked by atezolizumab. Human IL-2 ELISA showed that *hu*PD1 and chimeras with PEQ-containing ICDs (derived from monkey PD1 and dog PD1) inhibited IL-2 production by ∼60%, whereas significantly less inhibition of IL-2 (35-40%) was seen for PD1 chimeras with mouse or rat PD1 ICD (**fig. S9B**). Collectively, our data suggest that the pre-ITSM sequence can predict the signaling strength of PD1 orthologs and that rodent PD1 proteins have weaker activity due to the lack of a PEQ-like motif.

### Gradual functional attenuation of PD1 during rodent evolution

The weaker PD1 function suggested unique events and selection effects occurred during rodent evolution. To evaluate possible evolutionary scenarios underlying this phenomenon, we aligned the PD1 sequence from 107 mammalian species, including 19 rodents from all four major suborders (2 from Hystricomorpha, 1 from Sciuromorpha, 3 from Castorimorpha and 13 from Myomorpha). Based on the tree topology of mammal phylogeny, we conducted maximum likelihood inference of positive selection and relaxation on different combinations of tree branches in both the rodent and primate lineages (see **Methods**). At the basal branch (most recent common ancestor, MRCA) of all rodents, we found significant positive selection (*P* < 0.02 by BUSTED program) (*44*). The same is true for MRCA of primates (*P* < 0.003). Intriguingly, all branches within the rodent clade exhibited significant relaxation of selection pressure (*k* = 0.8, *P* < 0.05 by RELAX program (*45*), which was not found in primates (*k* = 1.0, *P* > 0.5) (**table S1**). These results indicate a unique lineage-specific relaxation of selection pressure on PD1 protein sequence during rodent evolution, suggesting gradual attenuation of PD1 function within the rodent clade. In contrast, for other inhibitory immunoreceptor genes such as BTLA, TIGIT, LAG3 and PVRIG, either positive selection or selection intensification were observed within the rodent clade (**table S1**).

To validate this potential trajectory of functional change, we next performed ancestral sequence reconstruction (ASR) of PD1 for seven key evolutionary time points in the tree (nodes with numerical IDs in **Fig. 5C**, predicated AA sequences are shown in **table S2**), following common practices of ASR (*46, 47*) (see **Methods**). For each node, we replaced the ICD sequence of *hu*PD1 with the reconstructed ancestral PD1 ICD, expressed the chimera in Jurkat cells, and examined its ability to suppress IL2 secretion upon coculture with SEE-pulsed *hu*PDL1-expressing Raji cells. We found that while the inhibitory capacities of the primate MRCA (node 172) and the shared MRCA of rodents and primates (node 170) were comparable to that of *hu*PD1, the inhibitory capacities of PD1 decreased along the evolution trajectory within the rodent lineage, from the MRCA of rodent and rabbit (node 194) to modern mice (**Fig. 5D**). Specifically, the transition of pre-ITSM PEQ to H at the MRCA branch of all rodents (node 195, 60-65 million years ago in Paleogene, right after the K-Pg boundary extinction event ∼66 million years ago) (*48, 49*) introduced a major decrease in PD1 inhibitory capacity. This was followed by additional attenuation between nodes 198 and 200 and between node 200 and modern mice. These sequential events indicate a gradual attenuation of PD1 function within the rodent lineage, suggesting a rodent-specific change of selection pressure during evolution.

### PD1 ICD-humanization severely impairs mouse T cell anti-tumor activity

We next asked if humanization of PD1 increased its immunosuppressive capacity in a mouse tumor model, using adoptively transferred tumor-specific TCR-transgenic CD8^+^ T cells expressing either *mo*PD1 or ICD-humanized PD1. We isolated *Pdcd1*^−/−^ P14 TCR transgenic CD8^+^ T cells recognizing H-2D^b^ (MHC-I)-restricted lymphocytic choriomeningitis virus (LCMV)-derived peptide antigen gp33-41. Using a retrovirus encoding an exogenous PD1 (exoPD1) and cell surface marker Thy1.1 (**Fig. 6A**), we reconstituted each of the three exoPD1 variants: *mo*PD1^WT^, *mo*PD1*^hu^*^ICD^ (whose entire ICD was humanized), and *mo*PD1^PEQ^ (whose pre-ITSM sequence [H] was replaced with the human version [PEQ]), together with the cell surface marker Thy1.1, during *in vitro* anti-*mo*CD3ε/anti-*mo*CD28 stimulation. The three versions of exoPD1 were expressed at comparable levels, as evidenced by flow cytometry (**Fig. 6B**). We then adoptively transferred the primed, gene-modified P14 CD8^+^ T cells into mice inoculated with B16.gp33 melanoma cells, which presented gp33-41 peptide via H-2D^b^. Tumors grew fastest in mice transferred with *mo*PD1*^hu^*^ICD^-expressing P14 cells (P14-*mo*PD1*^hu^*^ICD^), and slowest in mice transferred with *mo*PD1^WT^-expressing P14 cells (P14-*mo*PD1^WT^). Intermediate tumor growth was seen in mice transferred with *mo*PD1^PEQ^-expressing P14 cells (P14-*mo*PD1^PEQ^), and in control mice that had not received P14 transfer (**Fig. 6C**). These results provide *in vivo* evidence that ICD-humanization of *mo*PD1 increases its T cell suppressive capacity, at least partly due to the pre-ITSM PEQ motif. The mechanism by which *mo*PD1*^hu^*^ICD^-expressing P14 cells accelerated tumor growth compared to the no transfer control is unclear, but suggests that *hu*PD1 not only inhibits the anti-tumor activity of T cells, but also promotes tumor growth through other mechanisms. We confirmed these results using *Cas9*^+/-^ P14 T cells, in which we deleted the endogenous PD1 and re-expressed exogenous *mo*PD1^WT^, *mo*PD1*^hu^*^ICD^, and *mo*PD1^PEQ^ using a retrovirus plasmid encoding a *Pdcd1*-targeting sgRNA, the exoPD1 variant, and Thy1.1 (**fig. S10**). These data validated the stronger inhibitory activity of *hu*PD1^ICD^ and the significance of the pre-ITSM PEQ motif.

**Fig. 6.**
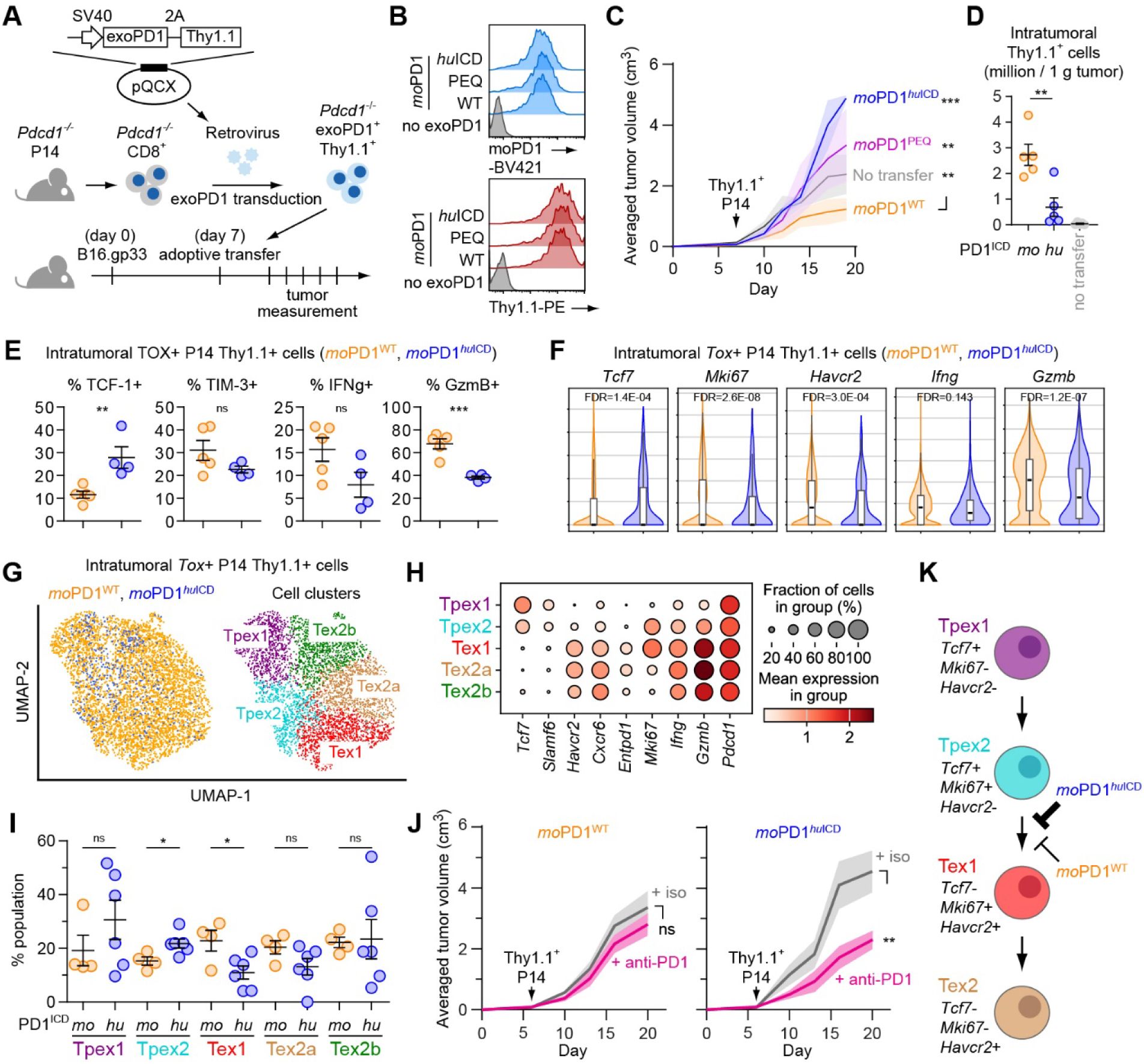
ICD humanization of *mo*PD1 inhibits precursor-to-terminal differentiation of *Tox*^+^ CD8^+^ T cells. (**A**) Schematic of an adoptive transfer experiment using *Pdcd1*^−/−^ P14 CD8^+^ T cells and B16.gp33 cells. *Pdcd1*^−/−^ P14 cells were retrovirally transduced with exoPD1 and Thy1.1, and adoptively transferred to mice bearing B16.gp33 melanoma. (**B**) Surface staining of indicated *mo*PD1 variants and Thy1.1 on P14 cells transferred to mice in **A**. (**C**) Tumor growth curves in mice received 1 million *Pdcd1*^−/−^ P14 cells expressing either *mo*PD1 or *mo*PD1*^hu^*^ICD^ (n = 2-4 mice). (**D**) Number of Thy1.1+ intratumoral CD8^+^ T cells (n = 5 tumors). (**E**) % TCF-1+, % TIM-3+, % IFNγ+, and % GzmB+ population within TOX+ intratumoral P14 cells containing either *mo*PD1 or *mo*PD1*^hu^*^ICD^, based on flow cytometry; n = 4-5 tumors. (**F**) Expressions of the indicated genes in *Tox*+ intratumoral P14 cells containing either *mo*PD1 or *mo*PD1*^hu^*^ICD^, based on scRNAseq. (**G**) UMAP showing the *Tox*+ intratumoral P14 cells containing either *mo*PD1 or *mo*PD1*^hu^*^ICD^ and their cell clusters. (**H**) Dot plot of indicated gene expressions in the cell subsets identified in **G**. (**I**) % population of the indicated subsets of intratumoral P14 cells containing either *mo*PD1 or *mo*PD1*^hu^*^ICD^; n = 4-6 tumors. (**J**) B16.gp33 tumor growth curves in mice that received 1.2 million P14 cells containing *mo*PD1^WT^ (left) or *mo*PD1*^hu^*^ICD^ (right), in response to treatment of either anti-PD1 (blue) or isotype control (magenta); n = 7 mice per group. (**K**) Model showing how PD1 humanization affects the precursor-to-terminal differentiation of *Tox*+ intratumoral CD8+ T cells. Data are mean ± SEM (**C**, **D**, **E**, **I**, and **J**). For violin-box plots (**F**), black bars are median, the boxes are 25% to 75% interquartile range, and the whiskers stand for minimum to maximum values excluding outliers. *P < 0.5; **P < 0.01; ***P < 0.001; ns, not significant; two-way ANOVA (**C** and **J**), student’s t-test (**D**, **E**, and **I**), or Wilcoxon signed-rank test (**F**).

### PD1 ICD-humanization accumulates the stem-like precursor exhausted T cells and lead to a stronger anti-PD1 response

To further investigate the cellular mechanism underlying the altered anti-tumor response associated with PD1 humanization, we characterized intratumoral T cells via flow cytometry and single-cell RNA sequencing (scRNA-seq). Flow cytometry showed that PD1-ICD humanization decreased the number of intratumoral P14 T cells (**Fig. 6D, fig. S11**) and their expression of granzyme B (GzmB), but did not alter their expression of IFNγ or the exhaustion marker TOX (**fig. S12A**). Consistent with the flow cytometry results, scRNAseq detected 8-fold more intratumoral P14 cells in mice receiving P14 (*mo*PD1) than in mice receiving P14 (*mo*PD1*^hu^*^ICD^) (**fig. S12B** and **C**). Gene expression analysis revealed that PD1-ICD humanization significantly reduced the expressions of *Gzmb* and *Mki67*, but not *Tox*, in intratumoral P14 T cells (**fig. S12D**). Uniform manifold approximation and projection (UMAP) identified five cell clusters, *Tox*^+^ clusters 0-2 and *Tox*^-^ clusters 3-4 (**fig. S12E**). PD1-ICD humanization did not alter the proportion of total *Tox*^+^ cell population (clusters 0-2), but increased the *Tox*^+^*Tcf7*^+^ subset (cluster 2) at the expense of *Tox*^+^*Tcf7*^-^ subsets (clusters 0 & 1) (**fig. S12F**). These data indicate that PD1-ICD humanization alters the differentiation within the *Tox*^+^ T cell population, a notion further supported by flow cytometry analysis at the protein level (**Fig. 6E**). Further scRNAseq analysis of the *Tox*^+^ population revealed that PD1-ICD humanization decreased *Mki67*, *Havcr2*, and *Gzmb* but increased *Tcf7* expression (**Fig. 6F**). UMAP of *Tox*^+^ cells identified five subsets (**Fig. 6G** and **H**): subset 1 expressed *Tcf7* and *Slamf6*, but lacked *Mki67*, consistent with a precursor-exhausted-1 T cell (Tpex1) signature; subset 2 expressed *Tcf7*, *Slamf6* and *Mki67*, consistent with a Tpex2 signature. Subsets 3-5 lost *Tcf7* but acquired *Havcr2* (Tim3), *Cxcr6* and *Entpd1* (CD39) expressions. Subset 3 maintained *Mki67* expression, consistent with a terminally-exhausted-1 T cell (Tex1) signature. Subset 4 and 5 were devoid of *Mki67*, consistent with a Tex2 signature (*50*), but differed in their cell cycle phases (**fig. S13**), and thus designated as Tex2a and Tex2b (**Fig. 6G** and **H**). PD1-ICD humanization reduced the Tex1 population, known to exhibit effector functions (*50*), while increasing the precursor-exhausted populations with less effector function (**Fig. 6I**). These data indicate that PD1-ICD humanization inhibited the Tpex2-to-Tex1 differentiation, which might have contributed to the lack of tumor control (**Fig. 6C** and **fig. S10D**). Indeed, anti-PD1 treatment elicited significant tumor control in mice receiving P14-*mo*PD1*^hu^*^ICD^, but not in mice receiving P14-*mo*PD1 (**Fig. 6J** and **fig. S14**), consistent with the relative abundance of Tpex under these two settings and prior reports that Tpex is the primary mediator for anti-PD(L)1 response (*51–54*). Altogether, our data suggest that *hu*PD1 more potently restricts the proliferation, effector function, and precursor-to-terminal transition of exhausted T cells compared to *mo*PD1 (**Fig. 6K**).

## DISCUSSION

Using quantitative assays, we document herein that *mo*PD1 uniquely has a weaker activity than all other PD1 orthologs tested. Specifically, we found that the weaker inhibitory activity of *mo*PD1 compared to *hu*PD1 mapped to both the ECD and ICD. With regards to intracellular signaling, *hu*PD1 and *mo*PD1 have been widely assumed to be functionally conserved, despite the merely 58% AA identity of their ICDs. This view was likely based on their identical ITSM, known as the major docking site for Shp2. However, the present study revealed that the ICD of *mo*PD1 is intrinsically less inhibitory than its human counterpart, and humanization of PD1 ICD disrupted the anti-tumor activity of CD8^+^ T cells.

Mechanistically, the lack of tumor control by P14 cells expressing ICD humanized PD1 stemmed from multiple factors, including the markedly decreased number of intratumoral T cells, lower effector function, and the altered T cell subset landscape (e.g. increased Tpex to Tex ratio). This notion is in line with the stronger anti-PD1 response in mice containing ICD-humanized PD1. These functional differences might contribute to the observed differential response to PD(L)1 inhibitors in mouse models and human patients, among other mechanisms. Given that this study relies on adoptively transferred T cells that express tumor-specific TCR, future studies are needed to investigate the impact of PD1 humanization on the anti-tumor immunity of endogenous T cells in the context of other tumor types.

Considering the role of PD1 in maintaining peripheral tolerance and preventing autoimmunity (*1, 4, 16, 18, 55*), our findings indicate that loss of PD1 activity could lead to a stronger autoimmune phenotype in humans than in mice. In fact, inherited PD1 deficiency in humans leads to death related to pneumonitis in two siblings at an early age (*18*). Moreover, anti-PD1 treatment in humans results in grade >3 adverse reactions in 10% of patients and any-grade toxicity in 20% of patients (*56, 57*). In contrast, mouse models of anti-PD1 do not recapitulate the adverse events seen in patients unless further bred onto an autoimmune susceptibility locus (*58*). In addition, PD1 deficiency in the C57BL/6 strain leads to lupus-like autoimmunity only when the mice were aged (>1 year) or when combined with the *<u>lpr</u>* mutation which promotes lymphoproliferation (*59*). Notably however, PD1 deficiency in the BALB/C strain leads to dilated cardiomyopathy and death as early as 5 weeks of age (*60*), suggesting that the strain background could regulate the function of PD1 in laboratory mice. While human vs mouse differences in PD1 deficiency penetrance and phenotype could stem from the strain, inbred nature, and specific pathogen free housing conditions of laboratory mice, our studies suggest the gene itself as another potential source of variation.

Humanized PD1 mouse models have been a valuable tool in testing human checkpoint inhibitors in pre-clinical settings, but they often express a PD1 chimera with only the ECD humanized, while the ICD of PD1 remained murine. Although such chimeric PD1 can respond to human PD1 inhibitors, the present study reveals intrinsic differences in the intracellular signaling of *hu*PD1 vs *mo*PD1, indicating that *mo*PD1 with only the ECD humanized cannot recapitulate *hu*PD1 function. Thus, mice knocked in with full length *hu*PD1 might be desirable for modeling human PD1 signaling and responses to anti-*hu*PD1 therapy.

The poorly conserved PD1 sequence across human and mouse is contrasted by the exquisite conservation of Shp2 sequence, but this appears to represent a common feature of many signaling pathways, with receptor showing lower evolutionary conservation and the intracellular effectors displaying higher conservation. Other examples include: CD28 and its effector PKCθ (*61*), which exhibit 68.8% and 95.0% sequence identities; BTLA and its effector Shp1 (*62*), which exhibit 49.2% and 94.5% sequence identities between human and mouse orthologs. Because intracellular effectors are often shared by multiple signaling pathways (*63*), their high degree of conservation might be required for the homeostasis of the cellular signaling network during evolution. Meanwhile, evolving the receptor sequence would offer a more specific and cost-effective means to reshape the functional outcomes of a particular signaling axis.

When conserved sequences often predict biological importance, our study emphasizes that sequence variations can inform new biological insights. Guided by this concept, we identify two novel motifs (PRS and PEQ), in conjunction with ITIM, that operate as modifiers of PD1 biochemistry and function. First and foremost, the PEQ motif, immediately upstream of ITSM in *hu*PD1 and altered to a single His in *mo*PD1, is primarily responsible for the stronger Shp2 binding activity of *hu*PD1. Indeed, the recently reported *hu*PD1-ITSM:Shp2 structure revealed hydrogen bonds between pre-ITSM residues and the SH2 domains of Shp2 (*64*). Second, the upstream PRS motif contributes significantly to PD1:Shp2 interactions. PRS might contain residues that directly interact with Shp2, or act by orienting ITIM and ITSM in a more optimal conformation to interact with Shp2. Third, the N-terminal phosphotyrosine motif ITIM exhibits species-specific biochemical features. Compared to *mo*PD1 ITIM, *hu*PD1 ITIM exhibited a greater affinity to Shp2 upon phosphorylation. These results might explain some of the discrepancies regarding the reported roles of ITIM in PD1 function. The divergence of these motifs confers functional plasticity in PD1 orthologs and offer avenues to engineering novel PD1 variants with desired functional properties.

The importance of the PEQ motif is highlighted by its perfect conservation in 131 out of 236 species we analyzed; PD1 orthologs in an additional 71 species contain PEQ-like motifs at the corresponding region. Rodent PD1 orthologs are interesting exceptions because this motif is substituted by a single His. The present study suggests that this replacement is the primary cause for the much weaker inhibitory activities of rodent PD1 compared to PD1 in other mammalian species. We further show that PD1 continued to undergo relaxation and weakening throughout rodent evolution, but we currently do not have data to explain why rodents would devalue and weaken PD1. Given that a weaker PD1 pathway might increase the strength and duration of immune responses and that this seemed to have occurred at the time of a major extinction event, it is possible that weakening PD1 (and thereby strengthening immune responses) afforded rodents with the ability to adapt to unique ecological niches and selective pressures from rodent specific pathogens.

## ACKNOWLEDGEMENTS

We thank J. Wilhelm and N. Stuurman for support in TIRF imaging, E. Griffis and P. Guo at the Nikon Imaging Center of UCSD for technical assistance in confocal imaging, P. Ordoukhanian at the Biophysics and Biochemistry Core of Scripps for performing BLI measurement, R. Ahmed and A. Sharpe for *Pdcd1*^−/−^ mice, K. Jepsen at the IGM Genomics Center of UCSD and L. Wang for technical assistance in scRNA-seq, A. Kamphorst, T. Wu and Y. Chen for technical advice, A.Y. Rudensky, Y. Liu and J. Chen for discussions, I. Mellman, and members of the Hui lab for critically reading the manuscript. Microscopy and image analysis were performed at the Nikon Imaging Center or the Hui lab at UCSD. This manuscript includes data generated at the UC San Diego IGM Genomics Center utilizing an Illumina NovaSeq X Plus that was purchased with funding from a National Institutes of Health SIG grant (#S10 OD026929). T. Masubuchi is supported by a postdoctoral fellowship from the Human Frontiers Science Program and JST PRESTO grant (JPMJPR22EB). This work was supported by R37 CA239072 from the National Institute of Health, a Searle Scholar Award from the Kinship Foundation, and a Pew Biomedical Scholar Award from the Pew Charitable Trusts to E. Hui. J. Bui was supported in part by The Hartwell Foundation.

## AUTHOR CONTRIBUTIONS

T.M., J.D.B. and E.H. designed the project and wrote the manuscript. J.D.B., J.P.M., N.M., and S.M.H. designed the adoptive transfer experiment. N.M. and S.M.H. established the retrovirus system to simultaneously delete endoPD1 via CRISPR and reconstitute exoPD1 in mouse primary T cells. G.A.W. cloned constructs for Figs. 4 and 5 and structural representation of Shp2. L.C. and Z.Z. designed and conducted sequence evolution analyses and computational reconstruction of ancestral sequences. J.D.B., T.M., N.M., C.C., and Y.Z. conducted *in vivo* experiments. J.Z. prepared human primary T cells expressing PD1 variants. T.M. performed all the other experiments and data analyses. E.H., J.D.B, L.-F.L. and X.C. obtained funding and supervised the project.

## DATA AND MATERIALS AVAILABILITY

All data are available in the main manuscript or the supplementary materials. Requests for materials should be addressed to E.H.

## Supplementary Materials for

### This PDF file includes

Materials and Methods

Figs. S1 to S14

Table S1 to S2

Captions for Movies S1 to S2

### Other Supplementary Materials for this manuscript include the following

Movies S1 to S2

## Materials and Methods

### Reagents

cDNA of rat PD1 (#RG80448-G), dog PD1 (#DG70109-G), and monkey PD1 (#KG90305-G) were purchased from Sino Biological. OVA_323-339_ peptide (#AS-27024) was purchased from Anaspec. Recombinant human IL-2 (#200-02) and recombinant mouse IFNγ (#315-05) were purchased from PeproTech. Staphylococcal enterotoxin E (SEE, #ET404) and staphylococcal enterotoxin B (SEB, #BT202) were purchased from Toxin Technology. RPMI-1640 media (#10-041-CM) was purchased from Corning. DMEM (#MT10017CV), MEM Non-Essential Amino Acids Solution (NEAA: #11140050), sodium pyruvate (#11360070), Alexa Fluor647 NHS ester (AF647 NHS ester: #A37573), Ni-NTA agarose (#88223), Zeba Spin Desalting Columns (#89890), dynabeads T-activator CD3/CD28 (#11132D), Polyethyleneimine (PEI: #NC1014330) and saponin (#558255) were purchased from Thermo Fisher Scientific. Fetal bovine serum (FBS: #FB-02) was purchased from Omega scientific. 1x Penicillin-Streptomycin (1x P/S: #SV30010) and Ficoll-Paque (#17544202) were purchased from Cytiva. SHP099 (#S8278) was purchased from Selleck Chemicals. FuGENE transfection reagent (#E2691) was purchased from Promega. Polybrene transfection reagent (#TR-1003-G) was purchased from EMD Milipore. Brefeldin A (#00-4506-51) and transcription factor staining kit (#00-5523-00) were purchased from eBioscience. Ghost DyeTM Red 780 (#18452) was purchased from Cell Signaling Technology. Hellmanex (#Z805939) was purchased from Sigma. Paraformaldehyde (PFA: #15714) was purchased from Electron Microscopy Sciences. Glutathione Agarose 4B (#G-250-50) was purchased from Gold Biotechnology. MojoSort human CD4 T Cell isolation kit (#480009), MojoSort human CD8 T cell isolation kit (#480012), MojoSort mouse CD8 T cell isolation kit (#480007), human IL-2 ELISA kit (#88702577), and mouse IL-2 ELISA kit (#431001) were purchased from Biolegend. Snap-Cell505 star (SC505: #S9103S) was purchased from NEB. DiD (#60014) was purchased from Biotium. Superdex 75 Increase column (#GE29-1487-21) and Superdex 200 Increase column (#GE28-9909-44) were purchased from GE Healthcare. Quantum PE MESF kit (#827) was purchased from Bangs laboratories. Glass-bottom 96-well plate (#P96-1.5H-N) was purchased from Cellvis. Collagenase (#LS004197) and soybean trypsin inhibitor (#LS003587) were purchased from Worthington Biochemical. eBioscience™ Foxp3 / Transcription Factor Staining Buffer Set (#00-5523-00) was purchased from Invitrogen. Chromium Next GEM Single Cell 3’ Reagent Kits v3.1 (Dual Index) with Feature Barcode Technology for Cell Surface Protein was purchased from 10x Genomics.

### Antibodies

Biotin anti-human CD3ε (clone OKT3, #317320) and biotin anti-mouse CD3ε (clone 145-2C11, #100303), APC anti-human CD69 (clone FN50, #310910), Alexa Fluor 700 anti-mouse CD8a (clone53-6.7, #100729), BV711 anti-mouse IFNγ (clone XMG.12, #505835), FITC anti-mouse CD8 (clone 16-10A1, #104705), BV785 anti-mouse CD8 (clone 53-6.7, #100750), PerCP/Cyanine5.5 anti-mouse CD8 (clone 53-6.7, #100734), PerCP/Cyanine5.5 anti-mouse CD45 (clone 30-F11, #103132), FITC anti-mouse Thy1.1 (clone OX-7, #202503), BV785 anti-mouse Tim3 (clone RMT3-23, #119725), FITC anti-mouse TCR Vα2 (clone B20.1, #127805), Alexa Fluor 647 anti-mouse Thy1.1 (clone OX-7, #202507), PE/Cyanine7 anti-human CD3 (clone UCHT1, #300419), APC anti-human CD4 (clone RPA-T4, #300552), PE anti-mouse CD8 (clone 16-10A1, #104707), and customized TotalSeq^TM^-anti-mouse Hashtag Antibodies devoid of anti-CD45 (#155831, #155833, #155835, #155839, #155841, #155843) were purchased from Biolegend. Anti-Shp2 (clone 79, #BDB610621) and BV605 anti-mouse Ly108 (clone 13G3, #745250) were purchased from BD Biosciences. Anti-mouse CD3ε (clone 145-2C11, #BE0001-1), anti-mouse CD28 (clone 37.51, #BE0015-1), InVivoMAb anti-mouse PD1 (clone J43, #BE0033-2), and InVivoMAb polyclonal Armenian hamster IgG (polyclonal, #BE0091) were purchased from Bioxcell. PE anti-human PD1 (clone MIH4, #12-9969-42) and PE-anti TOX (clone TXRX10, #12-6502-82) were purchased from Invitrogen. Pembrolizumab (#A2005) and atezolizumab (anti-PDL1, #A2004) were purchased from Selleck Chemicals. PE-Cy7 anti-mouse PD1 (clone J43, #25-9985-82) and PE anti-mouse Thy1.1 (clone HIS51, #12-0900-83) were purchased from eBioscience. Alexa Fluor 647 anti-TCF1/TCF7 (clone C63D9, #6709S) was purchased from Cell Signaling Technology. goat IgG fraction-anti-mouse IgG (#08670281) was purchased from MPbio. For TIRF imaging of Shp2 in **Figs. 4** and **5**, anti-Shp2 was labeled using AF647 NHS ester. The unreacted dye was removed via buffer exchange using a Zeba Spin Desalting Column (ThermoFisher), and eluted with HEPES buffer saline (50 mM HEPES-NaOH pH 7.5, 150 mM NaCl, 10% Glycerol), following manufacturer’s instructions. The purified antibody was stored at −80 °C until use.

### Lipids

1-palmitoyl-2-oleoyl-glycero-3-phosphocholine (POPC: #850457), 1,2-dioleoyl-sn-glycero-3-[(N-(5-amino-1-carboxypentyl)iminodiacetic acid)succinyl] (nickel salt) (DGS-NTA-Ni: #790404), 1,2-dipalmitoyl-sn-glycero-3-phosphoethanolamine-N-(lissamine rhodamine B sulfonyl) (ammonium salt) (Rhodamine-PE: #810158), 1,2-dipalmitoyl-sn-glycero-3-phosphoethanolamine-N-(biotinyl) (sodium salt) (Biotinyl-PE: #870285), and 1,2-dioleoyl-sn-glycero-3-phosphoethanolamine-N-[methoxy(polyethylene glycol)-5000] (ammonium salt) (PEG5k-PE: #880230) were purchased from Avanti Polar Lipids. N-(4,4-Difluoro-5,7-Dimethyl-4-Bora-3a,4a-Diaza-s-Indacene-3-Propionyl)-1,2-Dihexadecanoyl-sn-Glycero-3-Phosphoethanolamine (triethylammonium Salt) (Bodipy-PE: #D3800) was purchased from Thermo Fisher Scientific.

### Chimeric protein design

To construct *hu*PD1-based chimeras, *hu*PD1 ECD (aa24-170), TMD (aa171-191), or ICD (aa192-288) was replaced with the corresponding domain of *mo*PD1 (ECD: aa25-169, TMD: aa170-193, ICD: aa194-288), rat PD1 (ICD: aa194-287), dog PD1 (ICD: aa195-288), monkey PD1 (ICD: aa192-288), or reconstructed ancestral PD1 (**table S2**). *mo*PD1-based chimeras were designed in the same fashion using *mo*PD1 as a template. To construct GFP^ECD^-PD1^ICD^ chimeras, an N-terminal signal peptide of beta 2 microglobulin (MSRSVALAVLALLSLSGLEA), mGFP, human CD86 stalk-TMD region (aa236-271), and *hu*PD1 ICD or *mo*PD1 ICD were fused in sequence. To construct GFPNb-TM-TagBFP, GFPNb, an N-terminal signal peptide of beta 2 microglobulin (MSRSVALAVLALLSLSGLEA), the Fc region of human IgG1 (aa100-228), human CD86 stalk-TMD region, and TagBFP was fused in sequence. To construct PDL2-PDL1 chimeras, human PDL2 signal sequence (aa1-20), human or mouse PDL2 ECD (human: aa21-220, mouse: aa21-221), and human PDL1 TMD-ICD (aa238-290) were fused in sequence.

### Mice

Cas9 mouse were procured from Jackson Laboratory (B6(C)-Gt(ROSA)26Sorem1.1(CAG-cas9*,-EGFP)Rsky/J) and crossed with P14^+^ transgenic mice to generate *Cas9^+/-^* P14 mice. *Pdcd1*^−/−^ P14 mice were provided by Dr. Rafi Ahmed (Emory University). B6 mice were bred in-house. All animals were housed under specific pathogen-free conditions at the University of California San Diego (UCSD). All procedures were previously reviewed and approved by the UCSD IACUC under protocol S06201.

### Cell lines and cultures

Jurkat E6-1 cells (#TIB-152) and A20 cells (#TIB-208) were purchased from ATCC. HEK293T cells and Raji B cells were provided by Dr. Ronald Vale (University of California San Francisco). Phoenix cells were provided by Dr. Fuiboon Kai (University of California San Francisco). DO11.10 T cell hybridoma were provided by Dr. Philippa Marrack (National Jewish Health Center). HEK293T, Phoenix and B16.gp33 cells were maintained in DMEM media supplemented with 10% FBS and 1x P/S. Human-derived Jurkat cells and Raji B cells were maintained in RPMI media supplemented with 10% FBS and 1x P/S. Mouse-derived DO11.10 T cell hybridoma and A20 cells were maintained in RPMI media supplemented with 10% FBS, 1x P/S, and 50 µM β-mercaptoethanol. All cells were cultured in a 37 °C / 5% CO_2_ incubator.

### Cell line gene transduction and knock out

The gene of interest (GOI) was transduced into Jurkat cells and Raji B cells via lentivirus transduction, or into DO11.10 T cell hybridoma and A20 cells via retrovirus transduction. Lentivirus and retrovirus were prepared as described (*38*). For lentivirus production, GOI was cloned into pHR vector and co-transfected into HEK293T cells with envelope/packaging plasmids (pMD2.G and psPAX2) using PEI. For retrovirus production, GOI was cloned into pMIG II vector and transfected into Phoenix cells together with the pCL-Eco packaging plasmid using PEI. The virus supernatants were harvested 60-72 h post transfection, centrifuged to remove the contaminated cells, and mixed with target cells for transduction. To ensure similar expression of PD1 variants for quantitative comparisons, intrinsically high expressing constructs were transduced with diluted virus supernatant to reduce the expression level, whereas intrinsically low expressing constructs were re-infected to increase the expression level. The virus-containing media were replaced with fresh RPMI media 3-5 days post transduction. The cells expressing GOI were sorted using a FACS Aria flow cytometer (BD). Gene knock-out was performed utilizing CRISPR-Cas9 technique. Target cell lines were electroporated with pX330 vector using Gene Pulser Xcell (Bio-Rad). After the electroporation, the cells were cultured for 2-3 days and stained with corresponding antibodies to sort knocked-out cells using a FACS Aria flow cytometer (BD).

### Purification, culture, and transduction of human primary T cells

Human primary T cells were obtained from blood samples of healthy donors (San Diego Blood Bank) via a density gradient method and a magnetic beads purification. Briefly, human peripheral blood mononuclear cells (*hu*PBMC) were isolated from blood samples of healthy donors and enriched by Ficoll-Paque gradient centrifugation. The CD4^+^ or CD8^+^ T cells in the purified *hu*PMBCs were isolated using MojoSort Human CD4 T Cell Isolation Kit or MojoSort Human CD8 T Cell Isolation Kit (Biolegend) following the manufacturer’s instructions. The > 95% purity of the isolated T cells were confirmed via flowcytometric staining for CD3^+^CD4^+^ or CD3^+^CD8^+^ population. The purified T cells were pre-activated with Dynabeads T-Activator CD3/CD28 (Thermo Scientific) in the presence of 30 U/mL human IL-2 for two days before gene transduction. The pre-activated T cells were resuspended with lentivirus solution prepared as described above, and subjected to spinofection at 1000 x g for 90 min at 32 ⁰C. Human primary T cells were maintained in RPMI media supplemented with 10% FBS, 1x P/S, and 30 U/mL human IL-2 in a 37 °C / 5% CO_2_ incubator.

### PD1 CRISPR KO and reconstitution in mouse primary Cas9^+^ CD8^+^ T cells

For in vivo experiments in **fig. S10**, splenocytes from *Cas9^+/-^*P14 female mice were obtained by mashing spleens and lymph nodes through a 70-μm cell strainer (BD). Naïve CD8^+^ T cells were then enriched using Mojosort mouse CD8^+^ naïve T cell isolation kit (Biolegend). Purified naïve CD8^+^ T cells were resuspended in RPMI media supplemented with 5% FBS, 1 mM sodium pyruvate, 1x Penicillin-Streptomycin, 1x non-essential amino acid solution, and 55 μM β-mercaptoethanol, and activated using 1 µg/ml anti-mouse CD3ε, and 1 µg/ml anti-mouse CD28, incubated in a goat IgG fraction-anti-mouse IgG-coated plate at 37 °C / 5% CO_2_ for 24 h. Retrovirus pQCX plasmid encoding an exogenous *mo*PD1 variant (*mo*PD1^WT^, *mo*PD1^PEQ^, or *mo*PD1^huICD^), a P2A self-cleaving sequence, a Thy1.1 protein, and an sgRNA targeting endogenous *Pdcd1* were transfected to HEK293T cells using FuGENE transfection reagent following manufacturer’s instructions. Virus supernatants were harvested twice at 48 h and 72 h respectively, sterilized through a 0.45 µm filter (Millipore Sigma), incubated with the stimulated CD8^+^ T cells, and centrifuged at 1000 x g, 90 min at 32 °C for spinfection. After removal of the virus containing supernatant, cells were further cultured in the same RPMI medium supplemented with 100 U/mL human IL-2. 72-96 h later, Thy1.1^+^ cells were sorted and cultured in the same RPMI media supplemented with 100 U/mL human IL-2 for at least 48 h before the adoptive transfer. DNA sequences of the exogenous *mo*PD1 variants were silently mutated at the sgRNA-targeting site (ACACACGGCGCAATGACAG □ ATACCAGAAGGAACGATTC).

### Calculation of endo*PD1* KO scores for CD8^+^ T cells

DNA fragment of *Pdcd1* including the sgRNA-targeting site was amplified via polymerase chain reactions (PCR) using genomic DNA of *Cas9*^+/-^ CD8^+^ T cells as templates and using a primer pair (forward primer: TTCTGCATTTCAGAGGTCCCC, reverse primer: CCACCCACCCTACTTTGGC). The sequences of the amplicons were read via Sanger sequencing and subjected to the ICE program to calculate the % *Pdcd1*^−/−^ cells (KO scores) (*65*).

### PD1 reconstitution in mouse primary Pdcd1^−/−^ CD8^+^ T cells

For experiments in **Fig. 6**, and **figs. S11-14**, CD8^+^ T cells were isolated from *Pdcd1*^−/−^ P14 mice, stimulated for 24 h, and reconstituted with exogenous PD1 variant (*mo*PD1^WT^, *mo*PD1^PEQ^, or *mo*PD1^huICD^) as described in the previous section, except using a retrovirus lacking the endogenous *Pdcd1* targeting sgRNA. Specifically, the retrovirus was produced using a pQCX plasmid encoding an exogenous *mo*PD1 variant (*mo*PD1^WT^, *mo*PD1^PEQ^, or *mo*PD1^huICD^), a P2A self-cleaving sequence, and Thy1.1, without the sgRNA cassette.

### Adoptive transfer experiments

B6 mice were subcutaneously injected with 0.5-1 million B16.gp33 melanoma cells. 7 days after the B16.gp33 injection, P14 CD8^+^ T cells that were *in vitro* expanded for 7 days were transferred into the retro-orbital venous sinous in the B16.gp33-bearing mice under anesthesia (100 µL 1x PBS containing a mixture of pharmaceutical grade ketamine and xylazine). For measuring tumor growth, each tumor bearing mouse was transferred with 1 million (**Fig. 6C**), 1.2 million (**Fig. 6J** and **fig. S14**), or 0.5 million (**fig. S10**) of PD1-edited P14 CD8^+^ T cells. For measuring PD1 blockade effect on tumor growth, 100 µg anti-PD1 (clone J43) or isotype control was i.p. injected on day 3, 7, and 10 after T cell transfer. For measuring the absolute number of intratumoral Thy1.1^+^ CD8^+^ T cells shown in **Fig. 6D**, 2 million P14 CD8^+^ T cells were transferred to each tumor-bearing mouse and harvested from tumors 6 days after the transfer. For characterizing intratumoral Thy1.1^+^ CD8^+^ T cells, 1 million (**Fig. 6E** and **fig. S12A**) or 2 million (**Fig. 6F to I** and **fig. S12B to F**) P14 CD8^+^ T cells were transferred to each host mouse, and harvested from tumors 7 days after the transfer. Tumor sizes were measured twice or thrice per week by an investigator blinded to the identity of the animals. Sample size was determined by using at least seven animals per cohort when possible based on power calculations. Animals were randomized within each cage to receive one or the other type of T cells. If an animal had too much distress or the tumor was too large, this animal was sacked prior to flow cytometry experiments. Animals that died for any reason were excluded from the experiment. For characterization of intratumoral P14 Thy1.1^+^ T cells, tumors were harvested from mice on the indicated days, minced with razors and digested with 1 mg/mL collagenase type I and 1 mg/mL soybean trypsin inhibitor in 1x HBSS at 37 °C for 1 h, filtered through 70-μm cell strainer (BD), and resuspended in 1x PBS to prepare single-cell suspensions. The tumor single-cell suspension samples were subjected to percoll gradient centrifugation at 800 g for 10 min, after which CD8+ T cells were isolated using a CD8 beads selection kit (Biolegend).

### Flow cytometry analysis

For measuring protein expression on cell lines by antibody staining, cells were stained with the indicated antibodies following manufacturers’ instructions, washed with 1x PBS and resuspended in FACS buffer (PBS with 2% FBS), and subjected to flow cytometry analysis on an LSRFortessa X-20 cell analyzer (BD). For measuring protein expressions using mGFP or mCherry fluorescence, cells were directly resuspended in FACS buffer and subjected to flow cytometry. The absolute amount of PD1 surface expressions on human primary CD4^+^ T cells or on Jurkat cells was quantified using a Quantum PE MESF kit (Bangs laboratories) following the manufacturer’s instruction. Briefly, cells were stained with PE anti-human PD1 antibody and analyzed on an LSRFortessa X-20 cell analyzer together with the bead standards.

For characterizing intratumoral Thy1.1^+^ CD8^+^ T cells shown in **Fig. 6E** and **fig. S12A**, the isolated intratumoral CD8+ T cells were stimulated with 50 ng/mL phorbol 12-myristate-13-acetate (PMA) and 0.5 µg/mL ionomycin in the presence of 5 µg/mL Brefeldin A (BFA) and 5 µg/mL brefeldin A at 37 °C /5% CO2 for 4 h. After the stimulation, TILs were stained with Zombie Aqua Fixable Viability Dye (Biolegend, #423101) on ice for 15 min, with TruStain FcX antibody (biolegend, #101319) for 30 min, and subsequently with the antibodies for surface and intracellular protein markers using eBioscience™ Foxp3 / Transcription Factor Staining Buffer Set following the manufacturer’s instruction. Stained samples were analyzed on an LSRFortessa X-20 cell analyzer. The intratumoral Thy1.1^+^ CD8^+^ T cells were identified using the gates shown in **fig. S11**.

### Single cell RNA sequencing (scRNA-seq) sample preparation

Intratumoral CD8+ T cell sample isolated from tumor from each mouse was stained with Zombie Aqua Fixable Viability Dye (Biolegend, #423101) in 1x PBS on ice for 15 min, and subsequently with BV785 anti-moCD8, PE anti-Thy1.1 and FITC anti-TCRVα2 (P14 TCR) in 1x PBS containing 2% FBS on ice for 30 min. Live, CD8+, Thy1.1+, and TCRVα2+ cells in each sample were sorted on FACSAria into different tubes containing RPMI-1640 with 5% FBS. The sorted cell samples were distributed into two groups; cell samples isolated from mice that received *mo*PD1-expressing P14 cells, or cell samples isolated from mice that received *mo*PD1*^hu^*^ICD^-expressing P14 cells. Each cell sample in the same group was hash-tagged with a unique TotalSeq anti-mouse MHC-I antibody (Biolegend) on ice for 30 min, washed with 1x PBS containing 2% FBS. After the hash-tagging, the cell samples in the same group were pooled into one tube and diluted in 1x PBS containing 2% FBS to 1 million/mL. Approximately 20,000 and 2,000 cells were recovered in *mo*PD1 or *mo*PD1*^hu^*^ICD^ groups, respectively, all of which were subjected to library preparation. Both the gene expression (GEX) library and the hash-tag library of each group were prepared using a Chromium Next GEM Single Cell 3’ Reagent Kits v3.1 (Dual Index) with Feature Barcode Technology for Cell Surface Protein (10x Genomics) following manufacturer’s instructions. The prepared libraries were quantified and quality checked using TapeStation (Agilent). The GEX libraries or hash-tag libraries from each group were combined to generate pooled GEX library and pooled hash-tag library, respectively. The pooled libraries were sequenced using NovaSeq X Plus (Illumina) with a depth of 2 billion reads for the pooled GEX library and 200 million reads for the pooled hash-tag library.

### scRNA-seq data analysis

Alignments and count aggregation of gene expression and hash-tag reads were performed using the Cell Ranger Count v7.1.0 pipeline on 10x Cloud with the default settings. Gene expression reads or hash-tag reads were aligned to mouse genome (mm10-2020-A) and the hash-tag sequence list, respectively. The quality control report showed that 10209 or 1367 cells were detected for *mo*PD1 and *mo*PD1*^hu^*^ICD^ sample, respectively. The downstream analyses were performed using the Scanpy toolkit (*66*). Cells were initially quality-filtered based on the number of detected genes > 200. Genes were quality filtered based on the expressing cells > 3 and the read counts > 0. Cells were quality-filtered based on the percentage of mitochondrial reads < 5% to remove dead cells, the total counts of gene reads < 7500 to remove doublets, and the total counts of gene reads > 500 to remove low quality cells. After the filtering, the transcript counts were normalized by using scanpy.pp.normalize_total function with the default settings and transformed to a logarithmic scale by using the scanpy.pp.log1p function with the default settings. Cells were further filtered based on the expression of *Cd3e* > 0 and *Cd8a* > 0 to select CD8+ T cells, resulting in 7569 *mo*PD1-expressing cells and 952 *mo*PD1*^hu^*^ICD^-expressing cells.

For identifying the tumor origin of each cell, hash-tag reads were separated into two groups based on their PD1 ICD origins. The hash-tag reads in each group were processed using the scanpy.pp.pca function with n_comp=3, the scanpy.pp.neighbors function with n_neighbors=30, and the scanpy.tl.umap function to compute the PCA coordinates, the nearest neighbors distance matrix, and the UMAP. The cells were further clustered into subgroups using the scanpy.tl.leiden function with resolution=0.45 or 0.5 for *mo*PD1 or *mo*PD1huICD group, and the hash-tag labels were manually annotated to each cell cluster.

For clustering of total P14 cells based on the gene expression, PCA coordinates of total cells were computed using the scanpy.pp.pca function with n_comps=30, the nearest neighbors distance matrix was computed using the scanpy.pp.neighbors function with n_neighbors=50, and the UMAP was computed using the scanpy.tl.umap function. The cells were further clustered into subgroups using the scanpy.tl.leiden function with resolution=0.4. For analyzing *Tox*+ P14 cells, the gene expression and hash-tag matrix data were filtered based on the expression of *Tox* > 0, and the *Tox*+ cells were clustered using the same method for total cells except that the leiden resolution was set to 0.5.

For **Fig. 6F** and **fig. S12D**, the gene counts in each group were visualized using the scanpy.pl.violin function and the false discovery rate (FDR) was calculated using the scanpy.tl.rank_genes_groups function with the method=’wilcoxon’ setting. The dot plots in **Fig. 6H** and **fig. S12E** were generated using the scanpy.pl.dotplot function with the default settings. The % population in each cell cluster in **Fig. 6I** and **fig. S12F** was calculated through dividing the number of cells in each cluster by the total number of cells for each hash-tag sample. For computing the cell cycle phase shown in **fig. S13**, signature genes for cell cycle phase were derived from previous study (*67*), and the score of each cell cluster was calculated using the scanpy.tl.score_genes_cell_cycle function with the default settings.

### Proteins

Streptavidin (#S888) were purchased from Invitrogen. BSA (#A-420-1) was purchased from Goldbio. Recombinant protein *hu*PD1-His (#10377-H08H), *hu*PDL1-His (#10084-H08H), *mo*PD1-His (#50124-M08H), *mo*PDL1-His (#50010-M08H), *hu*ICAM-1 (#10346-H08H), and *mo*ICAM-1 (#50440-M08H) were purchased from SinoBiological. Recombinant protein *hu*PD1^ECD^-His (#PD1-H5221), *hu*PDL1^ECD^-*hu*Fc (#PD1-H5258), *hu*PDL2^ECD^-*hu*Fc (#PD2-H5251), *mo*PD1^ECD^-His (#PD1-M5228), *mo*PDL1^ECD^-*hu*Fc (#PD1-M5251), *mo*PDL2^ECD^-*hu*Fc (#PD2-M5254) were purchased from AcroBiosystems. His-tagged human Lck (aa3-509) was expressed in the Bac-to-Bac baculovirus system and purified as described (*42*). All His-tagged PD1^ICD^ were cloned into pET28A vector and expressed in BL21(DE3) strain of Escherichia coli. The expressed proteins were captured by Ni-NTA agarose (ThermoFisher), washed thrice with low imidazole buffer (50 mM HEPES-NaOH, pH 8.0, 150 mM NaCl, 30 mM Imidazole, 7 mM β-mercaptoethanol) and eluted with high imidazole buffer (50 mM HEPES-NaOH, pH 8.0, 150 mM NaCl, 500 mM imidazole, 50 mM HEPES-NaOH, pH 8.0, 150 mM NaCl, 7 mM β-mercaptoethanol). The eluted proteins were gel filtered using a Superdex 75 Increase column (GE Healthcare) in storage buffer (50 mM HEPES-NaOH, pH 7.5, 150 mM NaCl, 1 mM TCEP, 10% glycerol); monomeric fractions were pooled, snap frozen and stored at −80 °C until use. C-terminally SNAP-tag-fused Shp1^tSH2^ (aa4-215) or Shp2^tSH2^ (aa5-218) was each cloned into the pGEX-6P2 vector that encode an N-terminal GST-tag followed by a preScission protease cleavage site (LEVLFQGP). The GST-fused proteins were expressed in BL21(DE3) strain of Escherichia coli, purified using Glutathione Agarose 4B (Gold Biotechnology), and eluted with preScission buffer (50 mM HEPES-NaOH, pH 7.5, 150 mM NaCl, 0.5 mM TCEP, 20 units/mL 3C protease) to remove the GST-tag. The eluted proteins were further gel filtered using a Superdex 200 Increase column (GE Healthcare) in storage buffer (50 mM HEPES-NaOH, pH 7.5, 150 mM NaCl, 1 mM TCEP, 10% glycerol) and the monomeric fractions were pooled and stored at −80 °C until use. The SNAP-tagged SH2 proteins used in the liposome reconstitution assay were labeled with Snap-Cell505 star and purified using Zeba Spin Desalting Columns (ThermoFisher) following manufacturer’s instruction. The labeled proteins were snap-frozen and stored at −80 °C until use.

### Coculture assays

For measuring IL-2 secretion and CD69 expression in Jurkat:Raji coculture assays in **Figs. 1** and **2**, Raji B cells expressing *hu*PDL1 or *mo*PDL1 were pre-incubated with indicated concentration of SEE in the presence of indicated concentration of Atezolizumab at 37 °C for 30 min. The pre-incubated Raji B cells were mixed with Jurkat cells expressing indicated PD1 variant in a 96-well plate and centrifuged at 300 x g for 1 min to initiate Jurkat:Raji contact. After incubating the cell mixtures at 37 °C / 5% CO_2_ for 6 h or 24 h, the amount of IL-2 in the medium was quantified using Human IL-2 ELISA kit (Biolegend) or CD69 expression on Jurkat cells was measured by FACS, respectively. For measuring CD69 expression on Jurkat cells, Raji cells were stained with ViaFluor 405 Cell Proliferation Kit (Biotium) before SEE-pulse. The Jurkat:Raji mixtures were stained with APC anti-human CD69, then analyzed on an LSRFortessa X-20 (BD). 405-SE positive cells were excluded in FACS data to specifically measure CD69 expression on Jurkat cells.

For measuring IL-2 secretion in DO11.10:A20 coculture assay in **Figs. 1** and **2**, A20 cells expressing *hu*PDL1 or *mo*PDL1 were incubated with or without 120 µg/mL Atezolizumab at 37°C for 30 min, then mixed with indicated concentration of OVA_323-339_ in a 96-well plate. The treated A20 cells were further mixed with DO11.10 T cell hybridoma and centrifuged at 300 x g for 1 min. The cell mixture was incubated at 37 °C for 24 h. The amount of IL-2 in the culture medium was measured using a mouse IL-2 ELISA kit (Biolegend).

For measuring IL-2 secretion in human primary CD4^+^ T:Raji coculture assay in **Fig. 2**, *CD80^−/−^* Raji B cells or *CD80^−/−^GFPNb-TM-TagBFP^+^-*expressing Raji B cells were pre-incubated with 0.5 µg/mL SEB at 37 °C / 5% CO_2_ for 30 min. Human primary CD4^+^ T cells were washed twice with fresh RPMI medium containing 10% FBS to remove IL-2, mixed with SEB-loaded Raji B cells in a round-bottom 96-well plate, and centrifuged at 300 x g for 1 min to initiate cell contact. The cell mixture was then incubated at 37 °C / 5% CO_2_. 12 h later, the amount of secreted IL-2 was quantified using Human IL-2 ELISA kit (Biolegend).

For measuring the Shp2-dependent PD1 functions in **fig. S6**, the indicated human or mouse T cells were serum-starved for 24 h, incubated with 0 or 30 µM SHP099 in a serum-free RPMI media for 2 h, and further incubated with 0 or 80 µg/mL anti-PD1 blockade antibody (human: pembrolizumab, mouse: clone RMP1-14) for 30 min. To stimulate human T cells, *hu*PDL1-expressing *PTPN11*^−/−^ Raji cells were treated with 0.5 µg/mL SEB for 30 min, mixed with SHP099-treated human T cells in a serum-free RPMI, and incubated for 6 h. To stimulate mouse T cells, B16.gp33 cells were treated with 100 U/mL mouse IFNγ in the culture media for 24 h, washed with 1x PBS thrice, and incubated with SHP099-treated mouse T cells in a serum-free RPMI for 6 h. The final concentrations of SHP099 was 0 or 5 µM in the T:APC mixtures. Secreted IL-2 in the supernatant was measured using Human or Mouse IL-2 ELISA Kit (BioLegend). All incubations were performed at 37 °C / 5% CO_2_.

% inhibition on the T cell stimulation indicators, IL2 secretion and CD69 expression, exerted by PD1 variants were calculated using the following formula: 100% x (1 – [the readout value under no blockade condition] / [the readout value under each blockade concentration]).

For measuring the enrichment of PD1 on Jurkat:Raji contact surfaces in **Fig. 3**, Raji cells expressing mCherry-tagged *hu*PDL1 or *mo*PDL1 were incubated with 0.06 ng/mL SEE in the presence or absence of 120 µg/mL Atezolizumab at 37 °C for 30 min. The treated Raji B cells were mixed with Jurkat cells expressing indicated mGFP-tagged PD1 variants in a 96-well plate and centrifuged at 300 x g for 1 min to initiate cell contact. The cell mixtures were incubated at 37°C for 2 min, fixed with 4% PFA at room temperature (RT) for 15 min, washed with PBS containing 2% FBS, and transferred into a glass-bottom 96-well plate (Cellvis). Cell images were acquired using a confocal microscope (Nikon, Ti2-E).

### Liposome reconstitution assay

The liposome reconstitution assay was performed as described with modifications (*42*). Briefly, large unilamellar liposomes (lipid compositions: 89.7% POPC, 10% DGS-NTA-Ni, 0.3% Rhodamine-PE; diameter: 200 nm) prepared as described (*42*) were attached with 50 nM His-tagged Lck and indicated concentration of His-tagged PD1^ICD^, and further mixed with 50 nM SC505-labeled Shp2^tSH2^. The liposome-protein mixture was incubated at RT for 40 min with continuous monitoring the F.I. of SC505 to establish a baseline, then added with 1 mM ATP, after which the F.I. of SC505 was monitored for an additional 60 min. The F.I.s of SC505 at 30 min after ATP addition were used to measure the *K*_d_ of Shp2^tSH2^ binding to each PD1^ICD^ variant.

### LUV-SLB binding assay

SLBs were prepared as described, with modifications (*28*). A 96-well glass-bottom plate (Cellvis) was incubated with 2.5% Hellmanex at 50 °C overnight and washed extensively with ddH2O. Glass-bottom wells were etched with 6 N NaOH at 50 °C for 90 min and washed with ddH2O and Hepes-buffered saline (HBS: 50 mM HEPES-NaOH pH 7.5, 150 mM NaCl). Small unilamellar vesicles (lipid compositions: 95.9% POPC, 2% Biotinyl-PE, 2% DGS-NTA-Ni, 0.1% PEG5k-PE), prepared via a freeze-thaw method as described (*68*), were mixed with 1 μg/mL DiD, added to the washed wells containing HBS, and incubated at 50 °C for 90 min and further at room temperature for 30 min to form stable SLBs. Subsequently, the wells were rinsed with HBS containing 1 mg/mL BSA to block the SLBs, and incubated with 3 nM *hu*PDL1-His or *mo*PDL1-His at RT for 1 h before washed with HBS containing 1 mg/mL BSA. LUVs (lipid compositions: 89.7% POPC, 10% DGS-NTA-Ni, 0.3% Bodipy-PE) were prepared as described above. LUVs (total lipid concentration: 0.17 mM) were incubated with 0.83 nM *hu*PD1-His or *mo*PD1-His at RT for 1 h, allowing each LUV to capture about five PD1-His molecules. The PDL1-attached SLB were then incubated with PD1-attached LUVs (total lipid concentration: 70 μM) at RT for 30 min, and subjected to TIRF-M.

### Cell-SLB assay

SLBs were prepared as described above. After SLB formation, the wells were rinsed with PBS containing 1 mg/mL BSA to block the SLBs, then incubated with 1 mg/mL streptavidin, 3 nM *hu*ICAM1-His, and indicated concentrations of *hu*PDL1-His or *mo*PDL1-His at RT for 1 h. For preparing SLBs for DO11.10 T cell hybridoma, *hu*ICAM1-His was replaced with the same concentration of *mo*ICAM1-His. The SLBs were then rinsed with PBS containing 1 mg/mL BSA to remove unbound proteins, and further incubated with 2 µg/mL biotin anti-*hu*CD3ε at RT for 30 min. The functionalized SLBs were washed with 1x PBS containing 1 mg/mL BSA and 1x imaging buffer (20 mM HEPES-NaOH pH 7.5, 137 mM NaCl, 5 mM KCl, 1 mM CaCl_2_, 2 mM MgCl_2_, 0.7 mM Na_2_HPO_4_, 6 mM D-glucose) and incubated at 37 °C for 15 min before addition of cells. Cells were resuspended in 1x imaging buffer, loaded onto SLBs, incubated at 37 °C for 10 min. After incubation, cells were fixed with 2% PFA at RT for 10 min and washed with PBS containing 1 mg/mL BSA. The fixed cells were further permeabilized with 0.1% saponin at RT for 30 min. For observing endogenous Shp2, the fixed and permeabilized cells were incubated with anti-Shp2*AF647 at 4 °C overnight, washed with 1x PBS, and fixed with 4% PFA. TIRF images were acquired on a Nikon Eclipse Ti microscope equipped with a 100x Apo TIRF 1.49 NA objective lens, controlled by the Micro-Manager (*69*).

### Biolayer interferometry (BLI) assays

BLI assays were performed on Octet R8 (Sartorius) using anti-human IgG Fc Capture (AHC) biosensor. AHC sensors were pre-soaked in 1x PBS for 30 min, and incubated with 5 μg/mL *hu*PDL1^ECD^-*hu*Fc or *hu*PDL2^ECD^-*hu*Fc, or 10 μg/mL *mo*PDL1^ECD^-*hu*Fc or *mo*PDL2^ECD^-*hu*Fc diluted in Kinetic buffer (1x PBS with 0.02% Tween-20, 0.1% BSA, and 50 μM EDTA) for 300 sec. Analyte protein *hu*PD1^ECD^-His or *mo*PD1^ECD^-His were diluted in a two-fold series in Kinetic buffer (0.15625, 0.3125, 0.625, 1.25, 2.5, 5, 10 μM). Ligand-coated AHC sensors were dipped into diluted PD1 proteins for 180 sec for association, and subsequently dipped into Kinetic buffer for 180 sec for dissociation. The generated data were fitted to a 1:1 kinetic model using the Octet Analysis Studio (Sartorius, ver. 12.2.2.26) to estimate *K*_d_ values.

### Image analysis

All image analyses were conducted using Fiji (*70*). PD1 enrichment shown in **Fig. 3C** was calculated by dividing the PD1 F.I. within the Jurkat:Raji contact zone by the PD1 F.I. outside the contact zone. The cluster indices in **Fig. 3, E** and **F** were calculated by dividing the F.I. of clustered PD1 by the F.I. of total PD1 signal. For measuring the F.I. of clustered PD1, TIRF images were subjected to the “Subtract Background” function with the “Rolling ball radius” set as 20 pixels to define the PD1 microclusters. The whole F.I. of raw images were measured as the F.I. of total PD1. For calculating the Shp2 binding to PD1 microclusters shown in **Figs. 4** and **5**, the F.I.s of PD1 and anti-Shp2 on PD1 microclusters were used. The green (PD1), and far-red (anti-Shp2) or red (mCherry-Shp2) channels of TIRF images were subjected to the “Subtract Background” function with the same setting described above, and the binary, processed green channel images were used to define the positions of PD1 microclusters. The background-subtracted images were masked with the binary PD1 microcluster images, and the F.I.s of the masked green, far-red, and red channels were measured for PD1, anti-Shp2, or mCherry-Shp2 F.I.

### Data analysis

Data were shown as mean ±SD or ± SEM, as indicated in the figure legends. The number of replicates were indicated in the figure legends. Statistical significance was evaluated by two-tailed Student’s t test, One-way ANOVA, or Two-way ANOVA in GraphPad Prism 5 (*p < 0.05; **p < 0.01; ***p < 0.001; ****p < 0.0001) as indicated in the figure legends. Data with p > 0.05 are considered statistically non-significant (ns). All data fittings were conducted using GraphPad Prism 5.0. The binding curves in **Fig. 4H**, and **fig. S8B** were fit using the “On site – Specific binding” model. The dose-response curves in **Figs. 1B** and **3F** were fit using the “log(agonist) vs. response -- Variable slope” model.

### Phylogenies and orthologous coding sequences of genes

Phylogenies in **Fig. 5A**, the inference of positive selection and relaxation, and ancestral sequence reconstruction were obtained from TimeTree5 (*71*) by querying with the species needed. Branch lengths in **Fig. 5A** were obtained from TimeTree as well. The orthologous coding sequences of respective genes were obtained from OrthoMaM v12a (*72*) for all mammal species. Non-mammal orthologs of PD1 were obtained from NCBI Orthologs for Gene ID 5133. PD1 sequences of Xenopus_tropicalis (Accession #: A0A803JYL7), Xenopus_laevis (Accession #: A0A1L8G9Q9), Alligator_sinensis (Accession #: A0A3Q0GJP7), and Alligator_mississippiensis (Accession #: A0A151MMB5) were obtained from Uniprot. For obtaining the states of the PD1 pre-ITSM sequences in **Fig. 5A**, we first aligned all PD1 orthologs by linsi algorithm in MAFFT v7.310 (*73*) and then manually checked the validity of alignment and the states of pre-ITSM sequence. For (1) the inference of positive selection and relaxation in PD1 and related genes and (2) ancestral sequence reconstruction of PD1, the corresponding coding sequence alignment of the gene was first retrieved from OrthoMaM database, and then filtered as follows. We removed gap-rich sequences by simultaneously requiring the alignment to (1) contain no less than five rodent species and to (2) contain no sequence with more than 10% gaps. If the two requirements cannot be satisfied, we increased the gap proportion requirement by 5% increments in (2) until both criteria were met. We then removed the sites that contain only gaps in all remaining species in the alignment and used the derived alignment for downstream analyses.

### Statistical tests of positive selection and relaxation in sequence data

The existence of positive selection and relaxation in sequence evolution was tested by the BUSTED and the RELAX program in the HYPHY package v2.5.8 (*44, 45*). For each gene to be tested, the filtered alignment and the corresponding tree topology were used as input. In both BUSTED and RELAX, we assigned foreground branches in the tree according to the following two scenarios: (1) the MRCA branch of the clade of species (all rodents or all primates), (2) all branches in the clade (i.e. the subtree under the MRCA branch) excluding the MRCA branch. All other parameters were set as default in the analyses. BUSTED conducts likelihood ratio test on whether a proportion of the foreground branch(s) experienced positive selection (branch-site test). RELAX conducts likelihood ratio test on whether selection intensity k on the foreground branch(s) is the same (k = 1) as the background branches (all other branches by default). Intensification and relaxation of selection corresponds to the cases in which k increases (k > 1) or decreases (k < 1).

### Ancestral sequence reconstruction

We followed common practices to conduct the ASR analysis by using the computational pipeline Lazarus (retrieved from https://github.com/cxhernandez/project-lazarus) (*74*). We started from two 107-species filtered sequence alignments, containing respectively the amino acid and codon sequences for PD1. Together with the 107-species tree topology as input, we first inferred the best-fit evolutionary parameters (branch lengths and substitution model) separately for the amino acid sequence alignment and the codon sequence alignment by IQTREE v1.6.12, constraining the best-fit substitution matrix to be chosen from JTT, WAG or LG. We then ran the Lazarus pipeline separately using the amino acid sequence alignment and the codon sequence alignment, together with the trees containing respective branch lengths as input. The other Lazarus command-line parameters were set as “--model jones.dat --asrv 4 --alpha 1 --codeml --gapcorrect -- branch_lengths ‘fixed’”. The “--outgroup” was set as all species in the tree excluding rodents. Lazarus called PAML v4 (*75*) for maximum likelihood ancestral state inference and then assigned gaps at individual sites in the alignment when necessary according to maximum parsimony. As a result, we obtained an ASR for amino acid sequences and another ASR for codon sequences. We compared the reconstructed codon and amino acid states at each site. For sites with inconsistent amino acid and codon ASR states, when the codon states were not parsimonious in terms of their translated amino acid states, we modified the codon states according to the corresponding reconstructed amino acid states and maximum parsimony rules. For the pre-ITSM sequence, we first manually assigned the maximum parsimony amino acid states or gaps for the three sites at each node, and then assigned the codon states where a non-gap amino acid state was assigned, according to the Lazarus output codon ASR result. The reconstructed sequences were then used for downstream experiments.

---

**Fig. S1.**
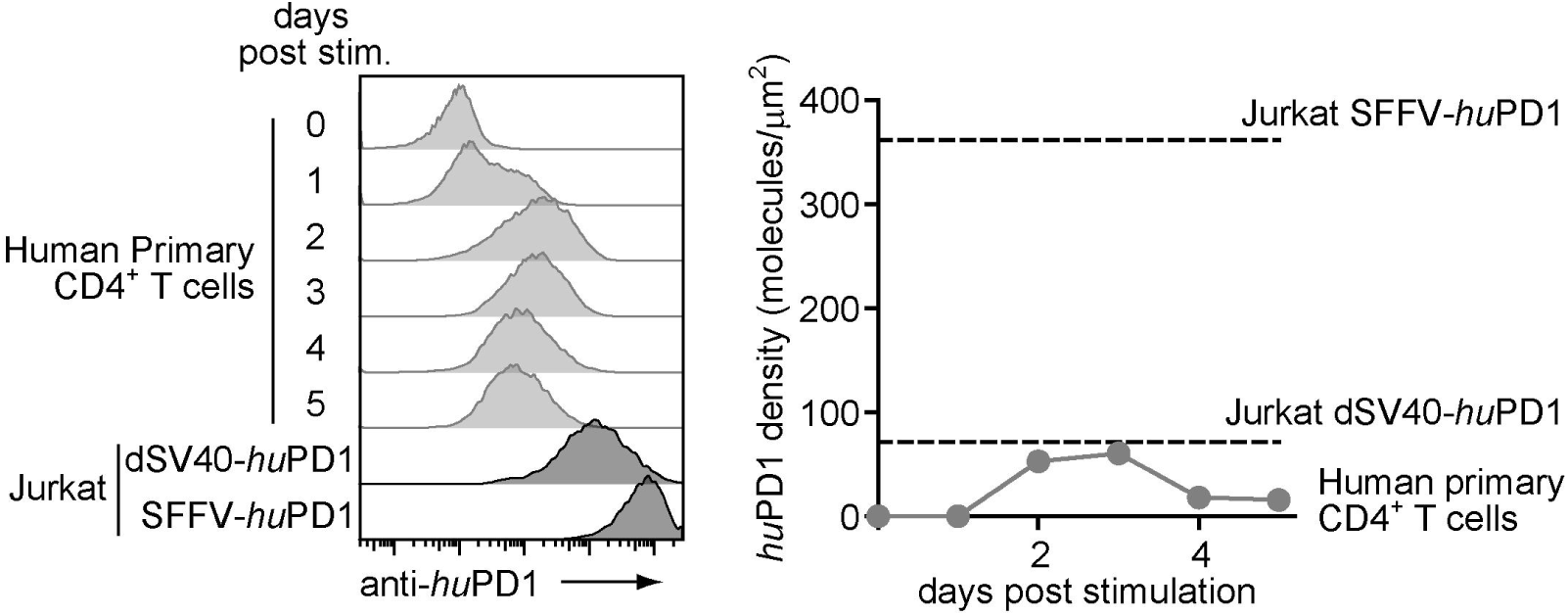
dSV40-driven PD1 expression level on Jurkat was similar to the endogenous PD1 expression level on human primary CD4^+^ T cells. Left, FACS histograms showing *hu*PD1 expressions on anti-CD3ε/anti-CD28 stimulated human primary CD4^+^ T cells and on stable Jurkat cells transduced with *hu*PD1 driven by either an dSV40 (weak) or an SFFV (strong) promoter. Right, PD1 densities on human primary CD4^+^ T cells plotted against duration of stimulation by anti-CD3/anti-CD28 beads. Dashed lines indicate the densities of SFFV- and dSV40-driven PD1 expressions on Jurkat cells. PD1 densities were calculated using molecular numbers quantified with a Quantum PE MESF kit and diameters of cells (human primary CD4^+^ T cell: 4.8 µm, Jurkat cells: 15 µm).

**Fig. S2.**
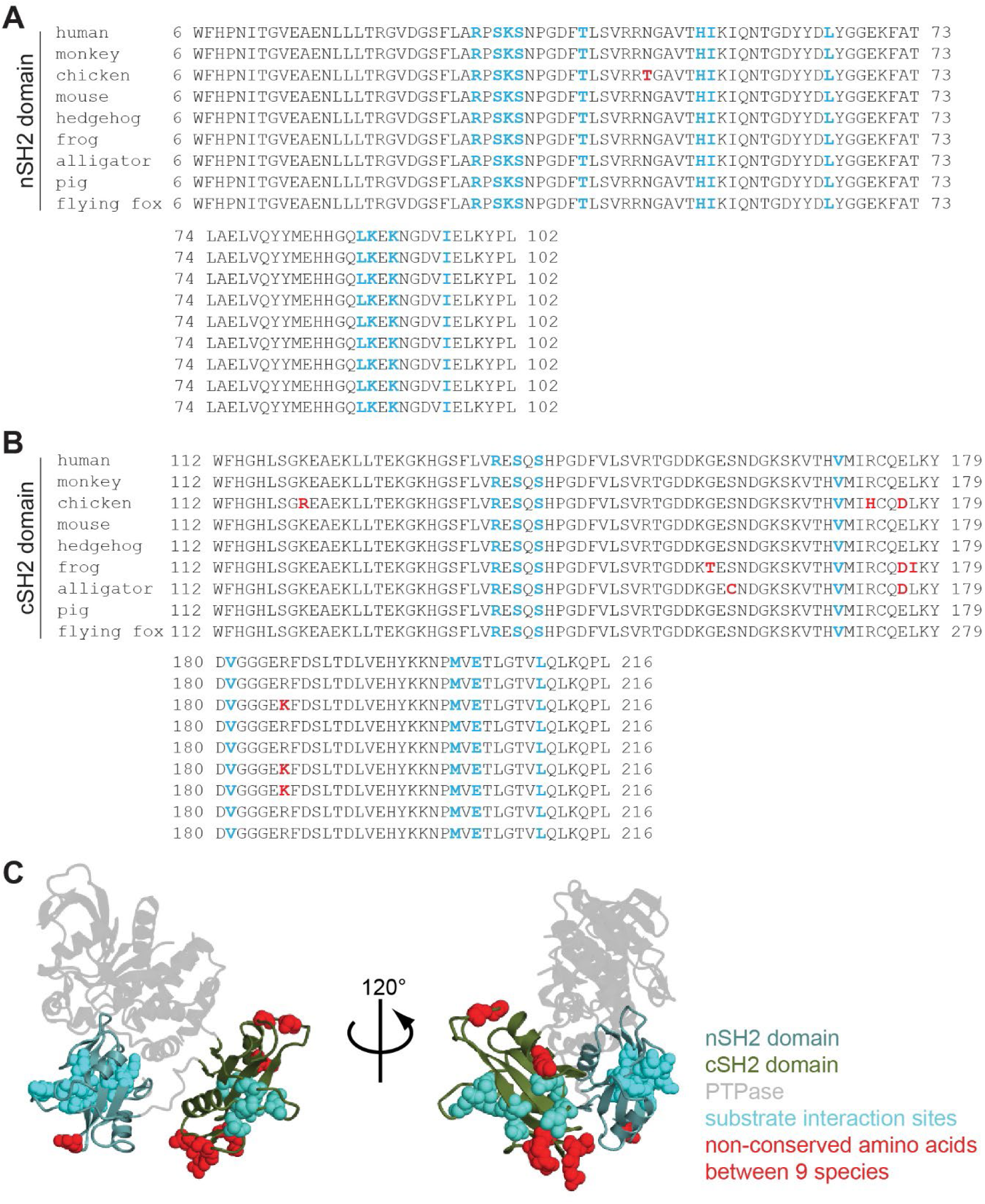
AA sequence alignments and structural representations of Shp2 in nine species. (**A**, **B**) Sequence alignment of Shp2 N-terminal SH2 (aa6-102) domain and C-terminal SH2 domain (aa112-216) in indicated nine species. AA residues involved in binding of phosphotyrosine motifs are shown in cyan, and non-conserved AAs are shown in red. (**C**) Ribbon structural representation of Shp2 (PDB: 4DGP). AA residues involved in phosphotyrosine binding (cyan) and non-conserved AA residues (red) are shown as space-filling models.

**Fig. S3.**
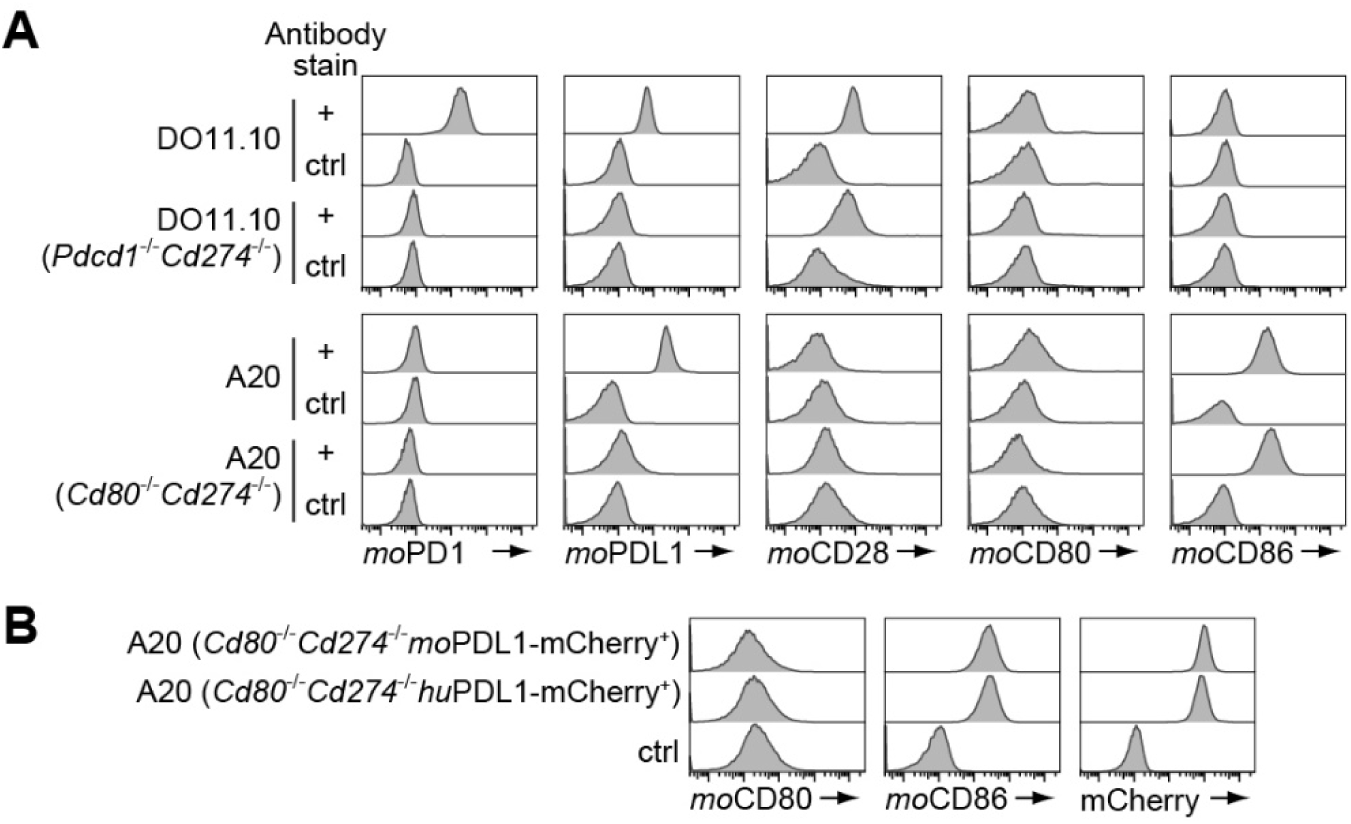
Expressions of endogenous moCD80, moCD86, *mo*PDL1 or lentivirally transduced mCherry-tagged *hu*PDL1 or *mo*PDL1 on A20 cells. **(A)** FACS histograms showing expressions of endogenous *mo*PD1, *mo*PDL1, *mo*CD28, *mo*CD80, and *mo*CD86 on DO11.10 T cell hybridoma and A20 cells before and after the KO of indicated genes. Isotype antibodies were used as controls. **(B)** FACS histograms showing *mo*CD80 and *mo*CD86 levels on A20 cells transduced with indicated mCherry-tagged PDL1 variants. Controls for *mo*CD80 and *mo*CD86 staining were generated with isotype antibodies. Control for PDL1-mCherry corresponded to untransduced *PDL1^−/−^* A20 cells.

**Fig. S4.**
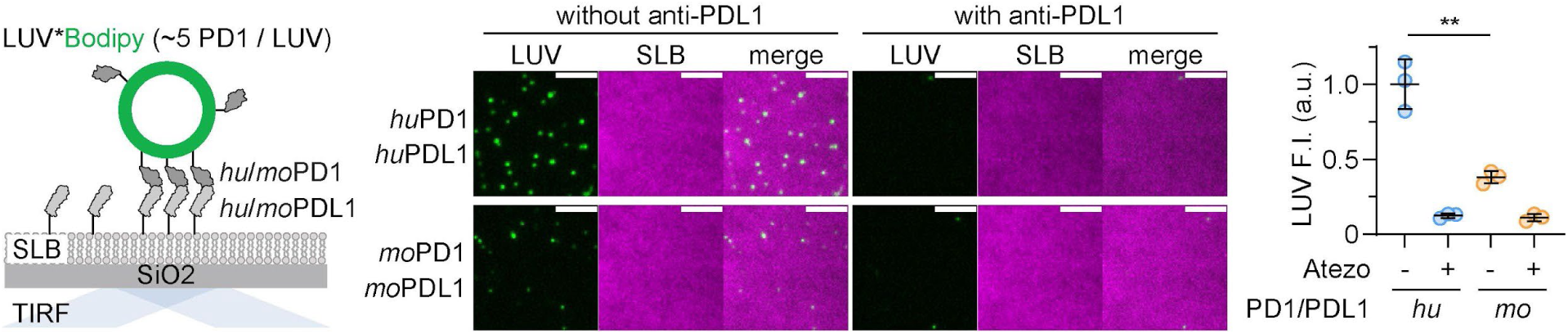
A LUV-SLB assay showing stronger *hu*PD1:PDL1 interaction compared to *mo*PD1:PDL1 interaction. Left, cartoon depicting a TIRF-M assay visualizing PD1^ECD^-attached, bodipy-labeled LUVs captured by PDL1^ECD^-attached SLB. Middle, representative TIRF images of LUVs (bodipy) and SLBs (DiD) in the presence or absence of anti-PDL1 blockade antibody (atezolizumab). Right, normalized F.I. of LUVs in the TIRF field under indicated conditions. Scale bars: 5 µm. Data are mean ± SD from three independent experiments. **P < 0.01; student’s t-test.

**Fig. S5.**
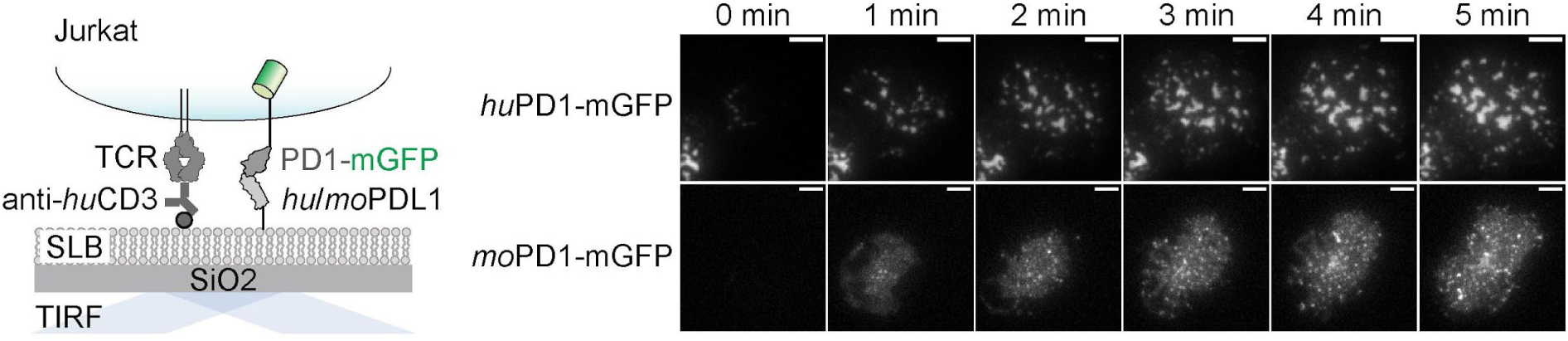
*hu*PD1 forms stronger microclusters than does *mo*PD1. **(A)** A cartoon depicting a cell-SLB assay using Jurkat expressing PD1-mGFP. **(B)** Time lapse TIRF images visualizing microclusters of indicated PD1 variants. Scale bars: 5 µm.

**Fig. S6.**
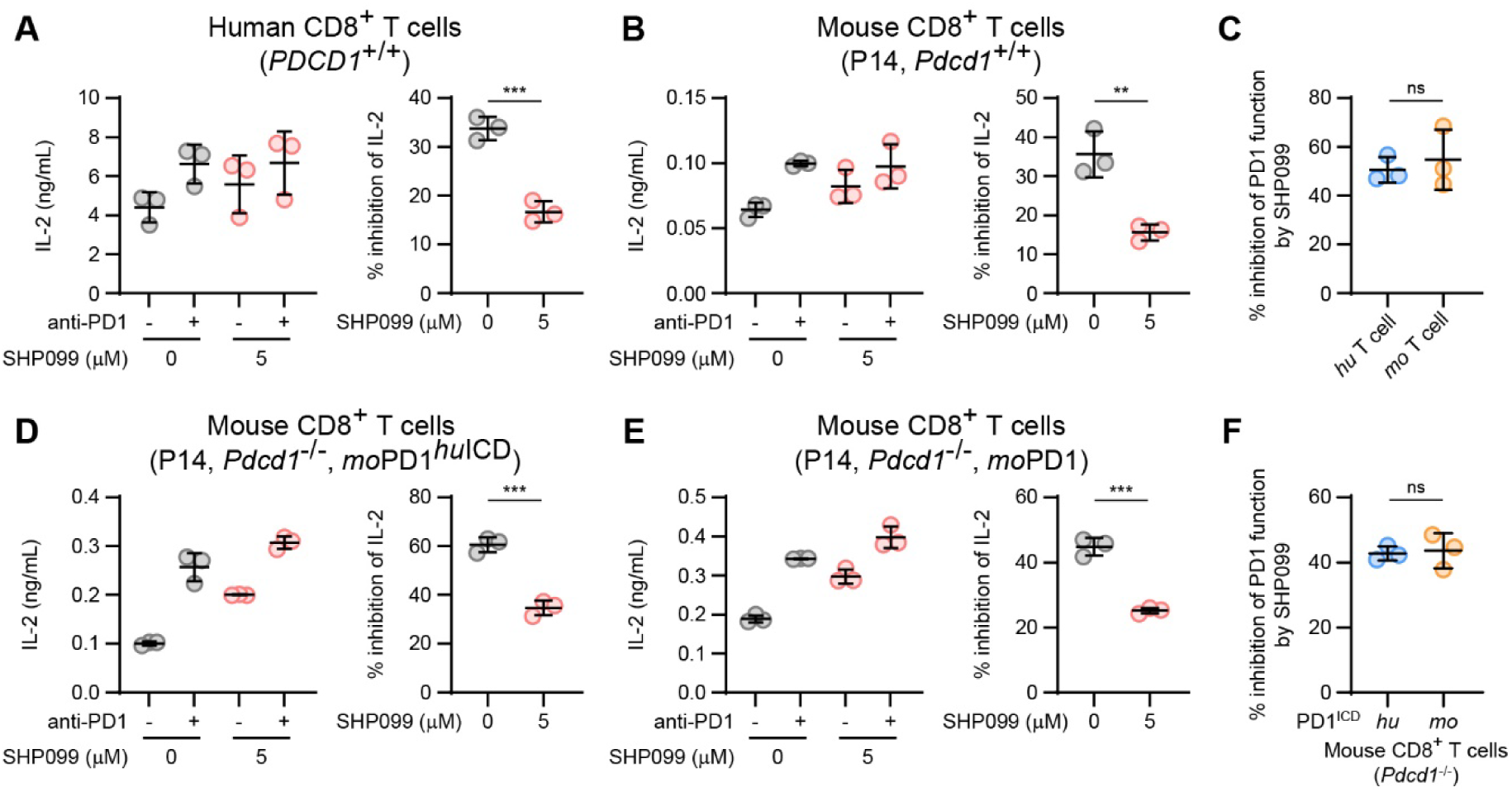
Shp2-dependencies of the suppressive functions of *hu*PD1 and *mo*PD1. **(A)** Left, IL-2 secretion from human CD8^+^ T cells stimulated by SEB-loaded *hu*PDL1-expressing Raji cells, with or without the presence of anti-PD1 blockade antibody (pembrolizumab), with or without 5 µM SHP099. Right, degree of PD1-mediated inhibition of IL-2 secretion with or without 5 µM SHP099, calculated based on data on the left. **(B)** Same as **A** except using mouse CD8^+^ T cells isolated from the splenocytes of P14 mice, and using anti-*mo*PD1 RMP1-14 to block PD1 signaling. **(C)** Degrees of SHP099 mediated inhibition of endogenous PD1 function in human and mouse CD8^+^ T cells calculated based on data from **A** and **B**. **(D)** Same as **B** except using *Pdcd1*^−/−^ P14 mouse CD8^+^ T cells engineered to express ICD-humanized *mo*PD1. **(E)** Same as **B** except using *Pdcd1*^−/−^ P14 mouse CD8^+^ T cells reconstituted with WT *mo*PD1. (F) Degrees of SHP099 mediated inhibition of exogenous PD1 function in *Pdcd1*^−/−^ P14 mouse CD8^+^ T cells calculated based on data from **D** and **E**. Data are mean ± SD from three independent experiments. **P<0.01, ***P < 0.001, ns, not significant; student’s t-test.

**Fig. S7.**
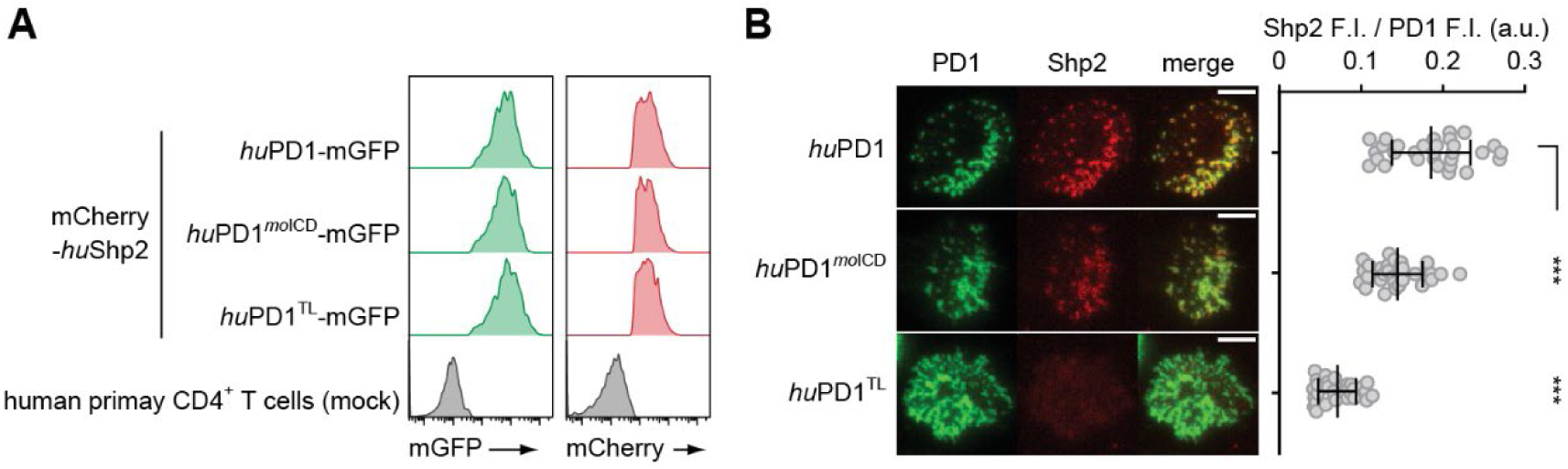
*hu*PD1^ICD^ more strongly recruited Shp2 than did *mo*PD1^ICD^ in human primary CD4^+^ T cells. **(A)** FACS histograms showing the expressions of indicated mGFP-tagged PD1 variant and mCherry-tagged huShp2 in human primary CD4^+^ T cells. T cells expressing both mCherry-huShp2 and the indicated PD1 variant were sorted and used in cell-SLB assays. **(B)** Left, representative TIRF images showing microclusters of indicated PD1-mGFP variants and mCherry-Shp2. Right, dot plots showing mCherry (Shp2) F.I. normalized to (mGFP) PD1 F.I. in TIRF images. Scale bars: 5 µm. Data are mean ± SD from n = 30 cells. ***P < 0.001; student’s t-test.

**Fig. S8.**
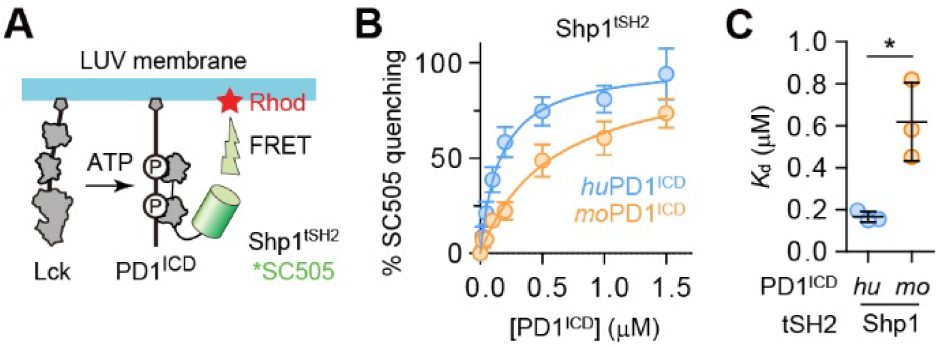
Shp1 more strongly binds to *hu*PD1 than to *mo*PD1. **(A)** Cartoon of a liposome reconstitution assay for measuring PD1^ICD^:Shp1^tSH2^ interaction. Same as **Fig. 4G** except using *hu*Shp1^tSH2^ rather than *hu*Shp2^tSH2^. **(B)** % ATP-triggered quenching of Shp1^tSH2^ fluorescence plotted against either [*hu*PD1-ICD] or [*mo*PD1-ICD]. **(C)** *K*_d_ values of PD1:Shp1^tSH2^ interaction determined via fitting the data in **B**. Data are mean ± SD from three independent experiments. *P < 0.05; student’s t-test.

**Fig. S9.**
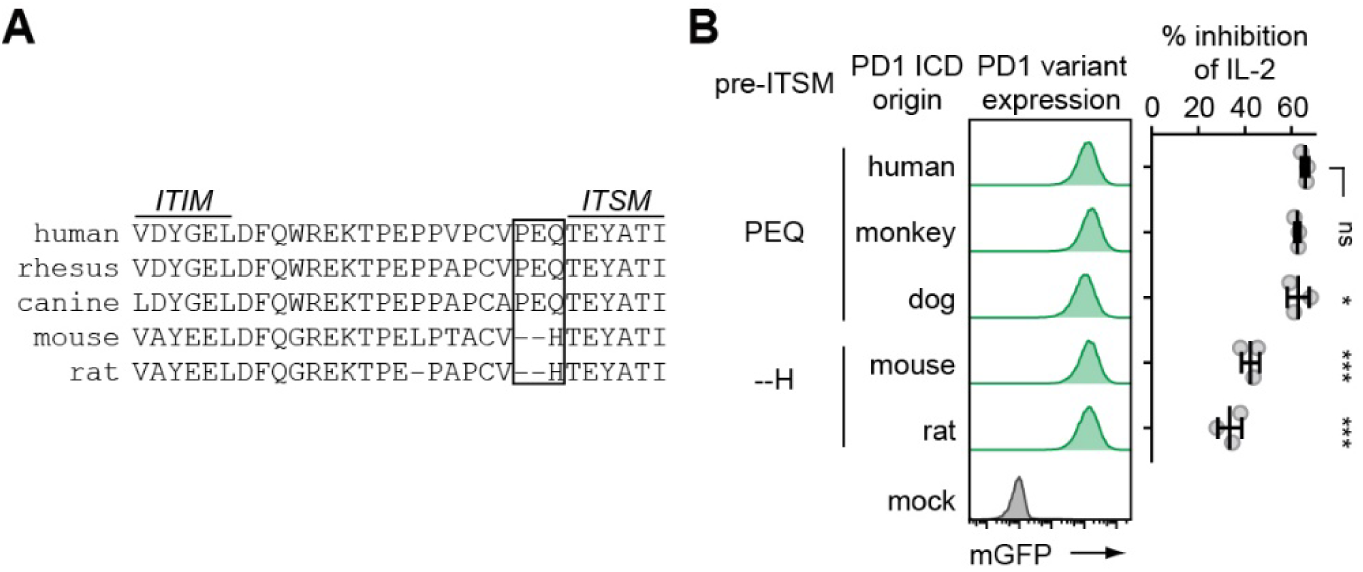
Rodents PD1 ICD are less inhibitory than PD1 ICD of other mammals. **(A)** AA sequence alignment of ITIM-ITSM region in PD1 of five selected species, with the pre-ITSM region annotated with a box. **(B)** Left, FACS histograms showing the expressions of mGFP-tagged *hu*PD1 variants that harbored the indicated PD1^ICD^ in five engineered Jurkat lines. Right, bar graphs showing % inhibition of IL-2 secretion mediated by the indicated *hu*PD1 variants. The five Jurkat lines were stimulated in parallel using SEE-pulsed, *hu*PDL1-expressing Raji cells, with or without Atezo (n = 3 independent experiments). Data are mean ± SD. *P < 0.05; ***P < 0.001; ns, not significant; student’s t-test.

**Fig. S10.**
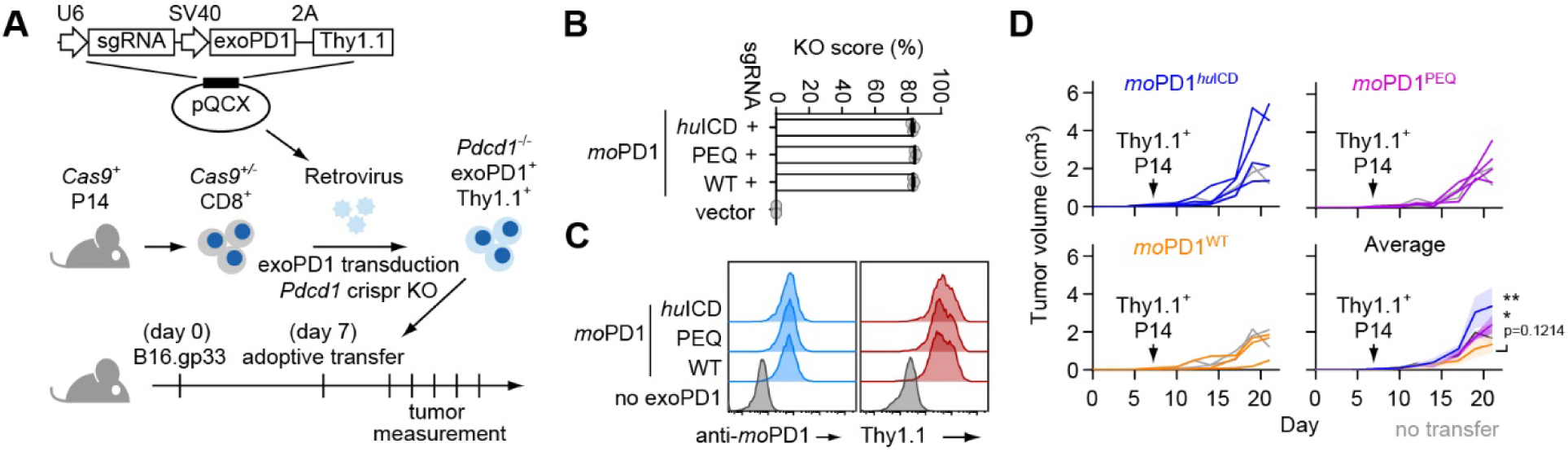
Cas9-mediated PD1KO and exoPD1 reconstitution revealed the stronger suppressive function of ICD humanized-PD1 for tumor-specific CD8 T cell function *in vivo*. **(A)** A diagram depicting an adoptive transfer experiment using *Cas9*^+/-^ P14 CD8^+^ T cells and B16.gp33 cells. *Cas9*^+/-^ P14 CD8^+^ T cells were retrovirally transduced with sgRNA targeting *Pdcd1*, exoPD1, and Thy1.1 to KO the *Pdcd1* and reconstitute with exoPD1. Thy1.1^+^ CD8^+^ T cells were sorted and adoptively transferred to mice bearing B16.gp33 melanoma. **(B)** *Pdcd1* KO scores of *Cas9*^+/-^ P14 CD8^+^ T cells calculated as described in Methods (n = 3 independent experiments). **(C)** Surface staining of indicated *mo*PD1 variants and Thy1.1 on T cells transferred to mice in **A**. **(D)** Tumor growth curves in individual mice received 0.5 million *Cas9*^+/-^ *Pdcd1*^−/−^ P14 CD8^+^ T cells expressing the indicated *mo*PD1 variant. Averaged tumor growth curves are shown in the bottom-right graph (n = 2-4 tumors). Data are mean ± SD (**B**) or SEM (**D**). *P < 0.05; **P < 0.01; ns, not significant; two-way ANOVA

**Fig. S11.**
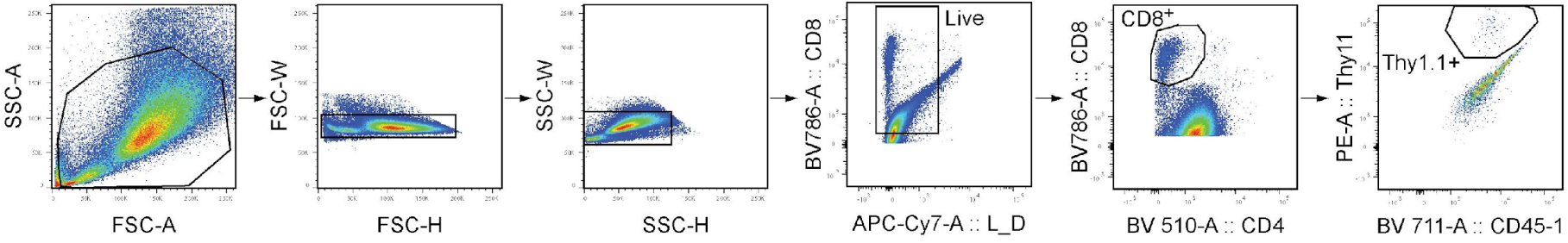
Gating strategy for Thy1.1^+^ CD8^+^ T cells in adoptive transfer experiments. Gating strategy for adoptively transferred Thy1.1^+^ CD8^+^ T cells in tumor samples.

**Fig. S12.**
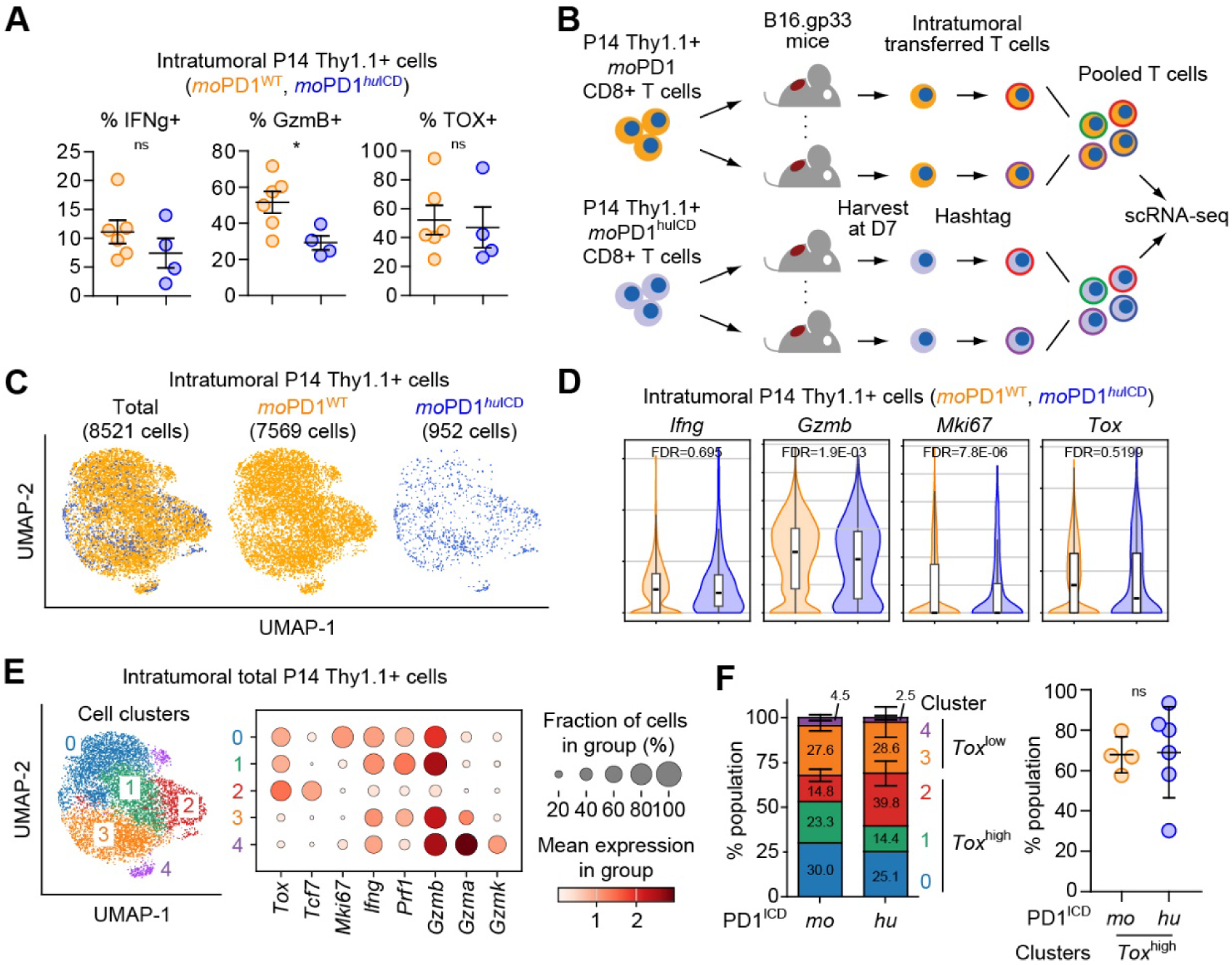
Characterization of intratumoral P14 CD8^+^ T cells expressing either *mo*PD1^WT^ or *mo*PD1^huICD^. **(A)** % IFNγ+, % GzmB+, and %TOX+ of the intratumoral P14 T cells expressing either *mo*PD1^WT^ or *mo*PD1*^hu^*^ICD^ measured by flow cytometry (n = 4-6 tumors). **(B)** Schematic showing scRNA-seq comparing intratumoral P14 Thy1.1+ CD8+ T cells expressing either *mo*PD1^WT^ or *mo*PD1*^hu^*^ICD^. Each mouse was adoptively transferred with 2 million P14 cells at 7 days after B16.gp33 inoculation. Another 7 days later, intratumoral P14 cells were purified, hash-tagged, and subjected to scRNA-seq. **(C)** UMAP showing the intratumoral P14 Thy1.1+ cells expressing *mo*PD1^WT^ (orange) or *hu*PD1*^hu^*^ICD^ (blue). **(D)** Expressions of the indicated transcripts in the intratumoral P14 Thy1.1+ cells expressing either *mo*PD1^WT^ or *mo*PD1*^hu^*^ICD^. **(E)** UMAP showing cell clusters (left) and the indicated gene expressions (right) of intratumoral P14 cells. Stem-like exhausted cells (cluster 2; *Tox*^high^, *Tcf7*^high^), Tex1 (cluster 0; *Tcf7*^low^, *Mki67*^high^), Tex2 (cluster 1; *Tcf7*^low^, *Mki67*^low^), *Prf1*-high effector cells (cluster 3; *Tox*^low^, *Pfr1*^high^, *Gzmb*^high^), and *Gzma/k*-high effector cells (cluster 4; *Tox*^low^, *Gzmb*^high^, *Gzma*^high^, *Gzmk*^high^) are annotated. (F) Left, % population of the indicated cell clusters in **E** for intratumoral P14 cells expressing either *mo*PD1 or *mo*PD1*^hu^*^ICD^. Right, summarized % population of *Tox*^high^ cell clusters shown in the left (n = 4-6 tumors). Data are mean ± SEM (**A** and **F**), or violin-box plot (**D**), in which the black bars are median, boxes are 25% to 75% interquartile range, whiskers are minimum to maximum values excluding outliers. *P < 0.05; ns, not significant; FDR, false discovery rate; student’s t-test (**A** and **F**), or Wilcoxon signed-rank test (**D**).

**Fig. S13.**
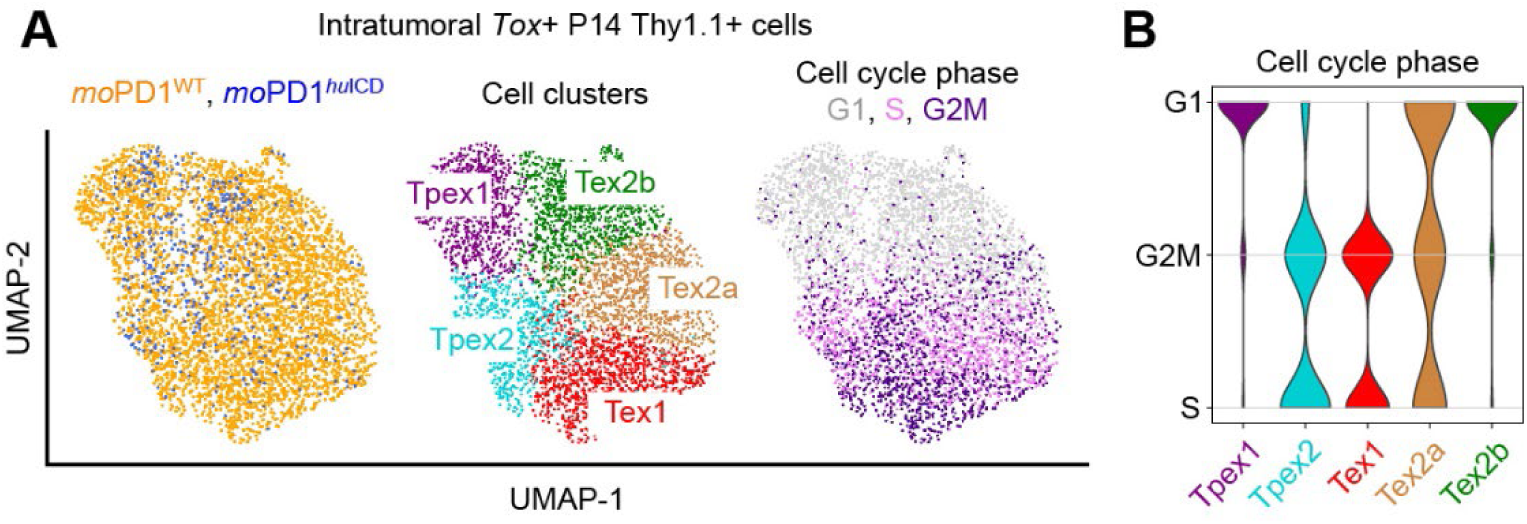
Differential cell cycle states of Tox+ intratumoral P14 cell subclusters. **(A)** UMAP showing the cell cycle phases of the five subclusters of *Tox*+ P14 cells expressing either *mo*PD1^WT^ or *mo*PD1^huICD^. **(B)** Violin plots showing the cell cycle phase of the five subclusters of *Tox*+ P14 cells.

**Fig. S14.**
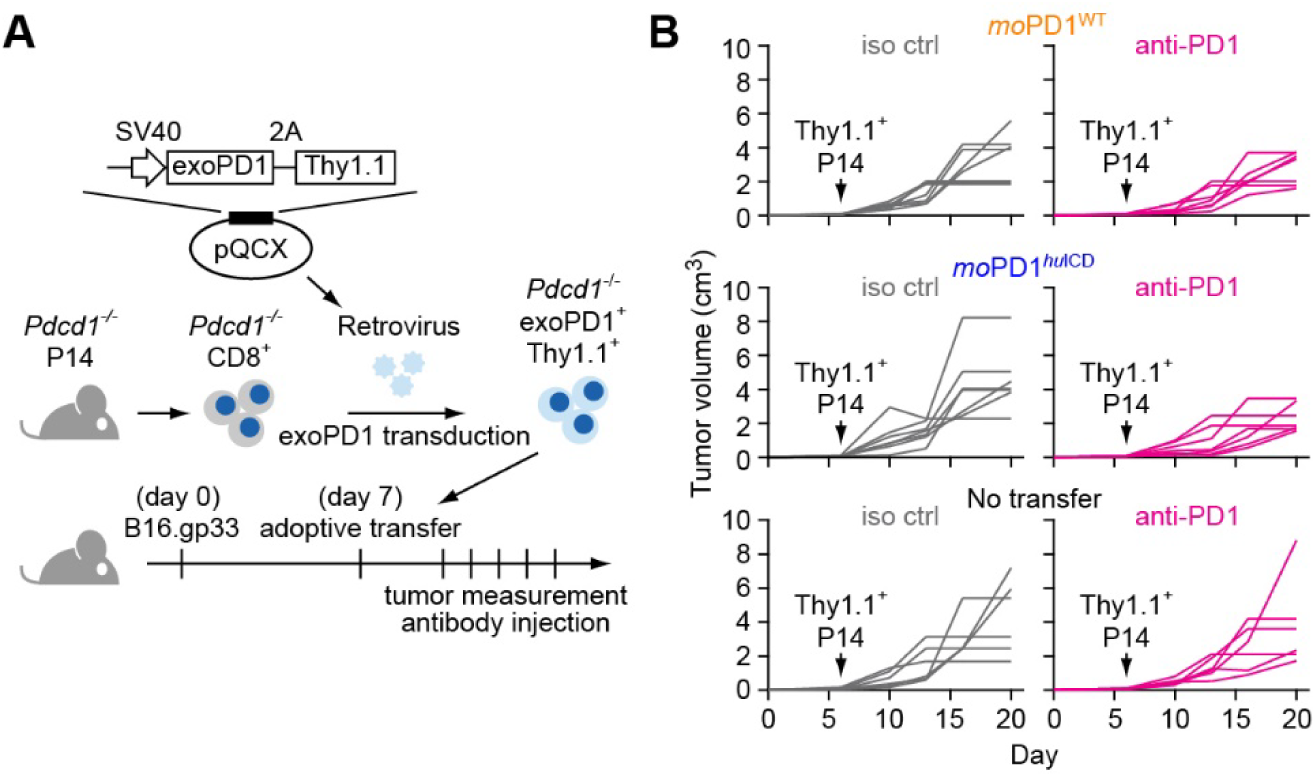
Tumor growth curves in individual mice that received P14 cells and/or PD1 blockade antibody. **(A)** Schematic of the adoptive transfer experiment. *Pdcd1*^−/−^ P14 cells were retrovirally transduced with exoPD1 and Thy1.1, and adoptively transferred to mice bearing B16.gp33 melanoma. The host mice then received anti-PD1 or isotype control on day 10, 14, and 17. **(B)** Tumor growth curves of individual mice that received 1.2 million *Pdcd1*^−/−^ P14 T cells expressing either *mo*PD1 or *mo*PD1*^hu^*^ICD^, and treated with either anti-PD1 or isotype control antibody.

**Table S1.**
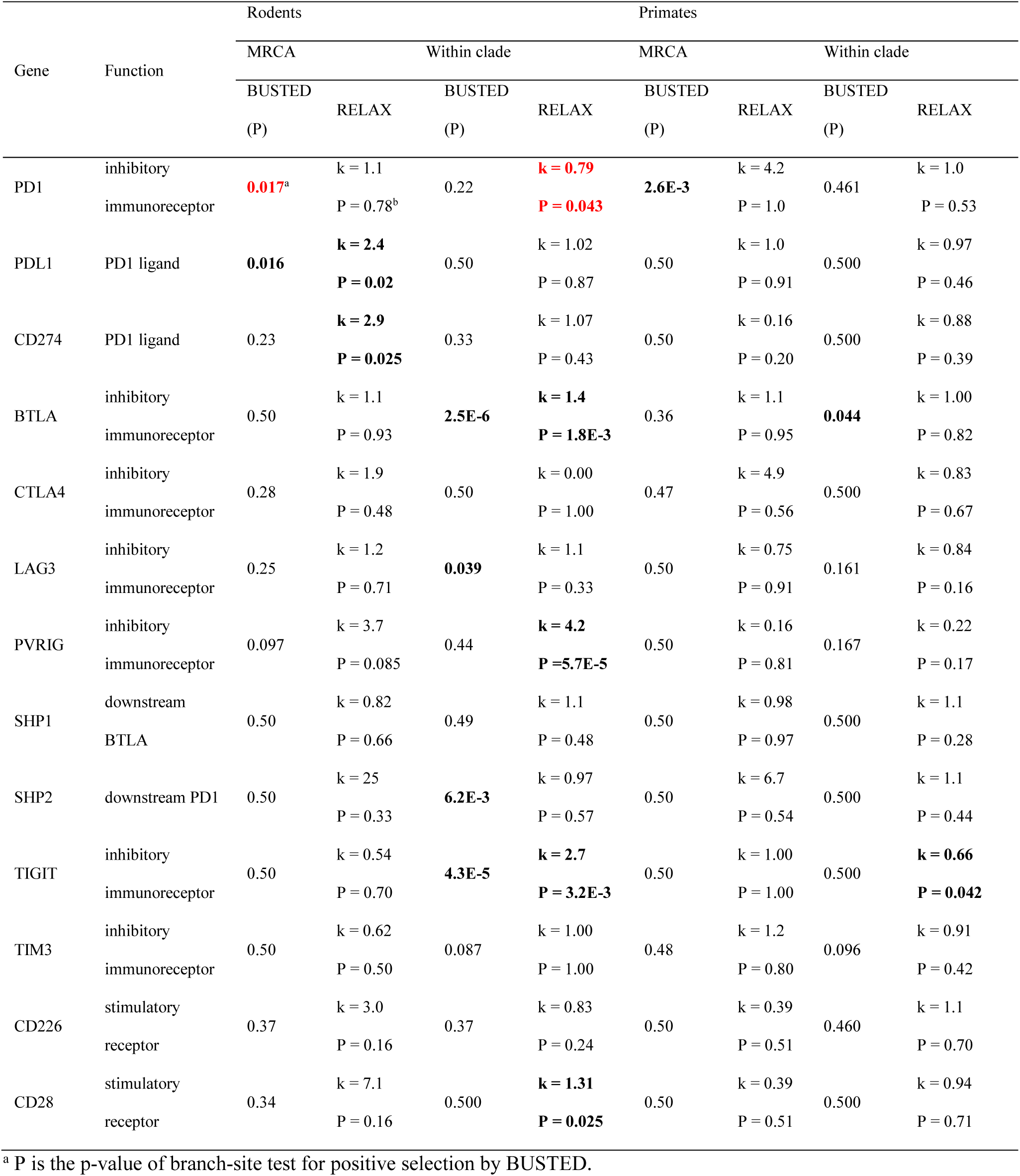
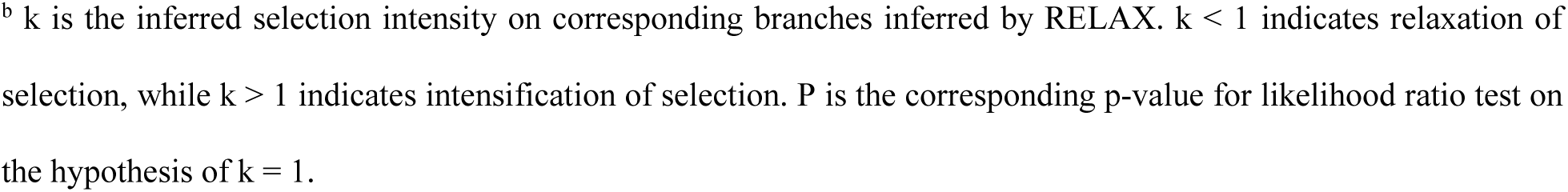
Test results of natural selection on sequences of PD1 and related genes in rodents and primates.

| Gene | Function | Rodents |  |  |  | Primates |  |  |  |
| --- | --- | --- | --- | --- | --- | --- | --- | --- | --- |
|  |  | MRCA |  | Within clade |  | MRCA |  | Within clade |  |
|  |  | BUSTED<br>(P) | RELAX<br>k | BUSTED<br>(P) | RELAX<br>k | BUSTED<br>(P) | RELAX<br>k | BUSTED<br>(P) | RELAX<br>k |
| PD1 | inhibitory | **0.017<sup>a</sup>** | k = 1.1 | 0.22 | **k = 0.79** | **2.6E-3** | k = 4.2 | 0.461 | k = 1.0 |
|  | immunoreceptor |  | P = 0.78<sup>b</sup> |  | **P = 0.043** |  | P = 1.0 |  | P = 0.53 |
| PDL1 | PD1 ligand | **0.016** | **k = 2.4** | 0.50 | k = 1.02 | 0.50 | k = 1.0 | 0.500 | k = 0.97 |
|  |  |  | **P = 0.02** |  | P = 0.87 |  | P = 0.91 |  | P = 0.46 |
| CD274 | PD1 ligand | 0.23 | **k = 2.9** | 0.33 | k = 1.07 | 0.50 | k = 0.16 | 0.500 | k = 0.88 |
|  |  |  | **P = 0.025** |  | P = 0.43 |  | P = 0.20 |  | P = 0.39 |
| BTLA | inhibitory | 0.50 | k = 1.1 | **2.5E-6** | **k = 1.4** | 0.36 | k = 1.1 | **0.044** | k = 1.00 |
|  | immunoreceptor |  | P = 0.93 |  | **P = 1.8E-3** |  | P = 0.95 |  | P = 0.82 |
| CTLA4 | inhibitory | 0.28 | k = 1.9 | 0.50 | k = 0.00 | 0.47 | k = 4.9 | 0.500 | k = 0.83 |
|  | immunoreceptor |  | P = 0.48 |  | P = 1.00 |  | P = 0.56 |  | P = 0.67 |
| LAG3 | inhibitory | 0.25 | k = 1.2 | **0.039** | k = 1.1 | 0.50 | k = 0.75 | 0.161 | k = 0.84 |
|  | immunoreceptor |  | P = 0.71 |  | P = 0.33 |  | P = 0.91 |  | P = 0.16 |
| PVRIG | inhibitory | 0.097 | k = 3.7 | 0.44 | **k = 4.2** | 0.50 | k = 0.16 | 0.167 | k = 0.22 |
|  | immunoreceptor |  | P = 0.085 |  | **P = 5.7E-5** |  | P = 0.81 |  | P = 0.17 |
| SHP1 | downstream | 0.50 | k = 0.82 | 0.49 | k = 1.1 | 0.50 | k = 0.98 | 0.500 | k = 1.1 |
|  | BTLA |  | P = 0.66 |  | P = 0.48 |  | P = 0.97 |  | P = 0.28 |
| SHP2 | downstream PD1 | 0.50 | k = 25 | **6.2E-3** | k = 0.97 | 0.50 | k = 6.7 | 0.500 | k = 1.1 |
|  |  |  | P = 0.33 |  | P = 0.57 |  | P = 0.54 |  | P = 0.44 |
| TIGIT | inhibitory | 0.50 | k = 0.54 | **4.3E-5** | **k = 2.7** | 0.50 | k = 1.00 | 0.500 | **k = 0.66** |
|  | immunoreceptor |  | P = 0.70 |  | **P = 3.2E-3** |  | P = 1.00 |  | **P = 0.042** |
| TIM3 | inhibitory | 0.50 | k = 0.62 | 0.087 | k = 1.00 | 0.48 | k = 1.2 | 0.096 | k = 0.91 |
|  | immunoreceptor |  | P = 0.50 |  | P = 1.00 |  | P = 0.80 |  | P = 0.42 |
| CD226 | stimulatory | 0.37 | k = 3.0 | 0.37 | k = 0.83 | 0.50 | k = 0.39 | 0.460 | k = 1.1 |
|  | receptor |  | P = 0.16 |  | P = 0.24 |  | P = 0.51 |  | P = 0.70 |
| CD28 | stimulatory | 0.34 | k = 7.1 | 0.500 | **k = 1.31** | 0.50 | k = 0.39 | 0.500 | k = 0.94 |
|  | receptor |  | P = 0.16 |  | **P = 0.025** |  | P = 0.51 |  | P = 0.71 |

<sup>a</sup> P is the p-value of branch-site test for positive selection by BUSTED.
<sup>b</sup> $k$ is the inferred selection intensity on corresponding branches inferred by RELAX. $k < 1$ indicates relaxation of selection, while $k > 1$ indicates intensification of selection. $P$ is the corresponding p-value for likelihood ratio test on the hypothesis of $k = 1$ .

**Table S2.**
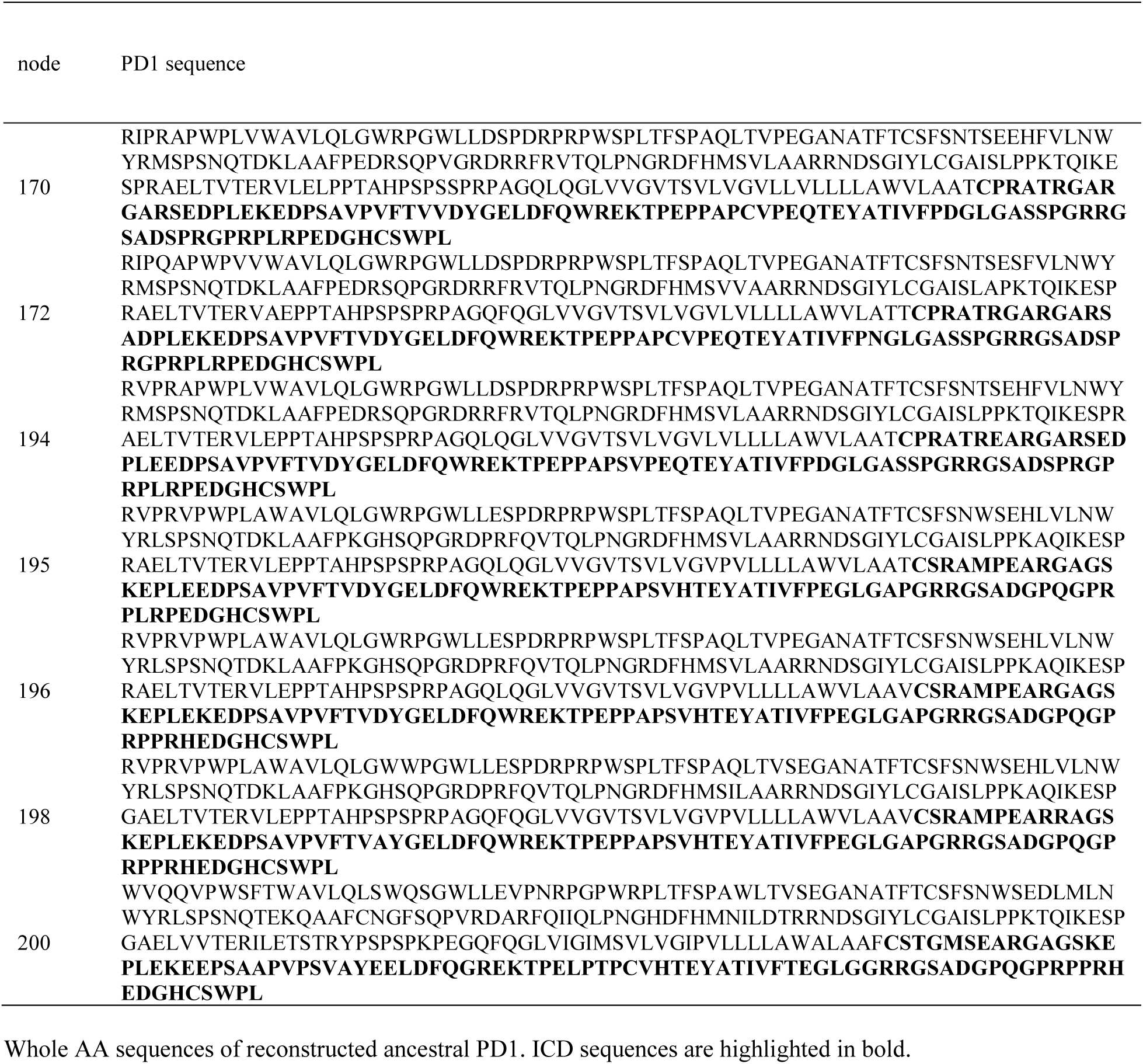
Reconstructed ancestral PD1 sequences.

## Movie S1

A movie showing microcluster formation of *hu*PD1 in a Jurkat cell upon SLB contact.

## Movie S2

A movie showing microcluster formation of *mo*PD1 in a Jurkat cell upon SLB contact.

## REFERENCES

1. L. M. Francisco, P. T. Sage, A. H. Sharpe, The PD-1 pathway in tolerance and autoimmunity. Immunol Rev 236, 219–242 (2010).

2. H. Dong et al., Tumor-associated B7-H1 promotes T-cell apoptosis: a potential mechanism of immune evasion. Nat Med 8, 793–800 (2002).

3. W. Zou, J. D. Wolchok, L. Chen, PD-L1 (B7-H1) and PD-1 pathway blockade for cancer therapy: Mechanisms, response biomarkers, and combinations. Sci Transl Med 8, 328rv324 (2016).

4. A. H. Sharpe, E. J. Wherry, R. Ahmed, G. J. Freeman, The function of programmed cell death 1 and its ligands in regulating autoimmunity and infection. Nat Immunol 8, 239–245 (2007).

5. L. Chen, Co-inhibitory molecules of the B7-CD28 family in the control of T-cell immunity. Nat Rev Immunol 4, 336–347 (2004).

6. D. R. Leach, M. F. Krummel, J. P. Allison, Enhancement of antitumor immunity by CTLA-4 blockade. Science 271, 1734–1736 (1996).

7. J. Mestas, C. C. Hughes, Of mice and not men: differences between mouse and human immunology. J Immunol 172, 2731–2738 (2004).

8. L. K. Beura et al., Normalizing the environment recapitulates adult human immune traits in laboratory mice. Nature 532, 512–516 (2016).

9. K. Medetgul-Ernar, M. M. Davis, Standing on the shoulders of mice. Immunity 55, 1343–1353 (2022).

10. M. F. Sanmamed, C. Chester, I. Melero, H. Kohrt, Defining the optimal murine models to investigate immune checkpoint blockers and their combination with other immunotherapies. Ann Oncol 27, 1190–1198 (2016).

11. R. Goodwin, G. Giaccone, H. Calvert, M. Lobbezoo, E. A. Eisenhauer, Targeted agents: how to select the winners in preclinical and early clinical studies? Eur J Cancer 48, 170–178 (2012).

12. A. Gunjur et al., Poor correlation between preclinical and patient efficacy data for tumor targeted monotherapies in glioblastoma: the results of a systematic review. J Neurooncol 159, 539–549 (2022).

13. J. T. Atkins et al., Pre-clinical animal models are poor predictors of human toxicities in phase 1 oncology clinical trials. Br J Cancer 123, 1496–1501 (2020).

14. N. L. Bayless et al., Development of preclinical and clinical models for immune-related adverse events following checkpoint immunotherapy: a perspective from SITC and AACR. J Immunother Cancer 9, (2021).

15. Y. Ishida, Y. Agata, K. Shibahara, T. Honjo, Induced expression of PD-1, a novel member of the immunoglobulin gene superfamily, upon programmed cell death. EMBO J 11, 3887–3895 (1992).

16. G. J. Freeman et al., Engagement of the PD-1 immunoinhibitory receptor by a novel B7 family member leads to negative regulation of lymphocyte activation. J Exp Med 192, 1027–1034 (2000).

17. D. L. Barber et al., Restoring function in exhausted CD8 T cells during chronic viral infection. Nature 439, 682–687 (2006).

18. M. Ogishi et al., Inherited PD-1 deficiency underlies tuberculosis and autoimmunity in a child. Nat Med 27, 1646–1654 (2021).

19. S. L. Topalian, C. G. Drake, D. M. Pardoll, Immune checkpoint blockade: a common denominator approach to cancer therapy. Cancer Cell 27, 450–461 (2015).

20. D. M. Pardoll, The blockade of immune checkpoints in cancer immunotherapy. Nat Rev Cancer 12, 252–264 (2012).

21. P. Sharma, S. Hu-Lieskovan, J. A. Wargo, A. Ribas, Primary, Adaptive, and Acquired Resistance to Cancer Immunotherapy. Cell 168, 707–723 (2017).

22. P. Sharma, J. P. Allison, Immune checkpoint targeting in cancer therapy: toward combination strategies with curative potential. Cell 161, 205–214 (2015).

23. A. Ribas, J. D. Wolchok, Cancer immunotherapy using checkpoint blockade. Science 359, 1350–1355 (2018).

24. G. J. Freeman, Structures of PD-1 with its ligands: sideways and dancing cheek to cheek. Proc Natl Acad Sci U S A 105, 10275–10276 (2008).

25. E. Lazar-Molnar et al., Crystal structure of the complex between programmed death-1 (PD-1) and its ligand PD-L2. Proc Natl Acad Sci U S A 105, 10483–10488 (2008).

26. D. Y. Lin et al., The PD-1/PD-L1 complex resembles the antigen-binding Fv domains of antibodies and T cell receptors. Proc Natl Acad Sci U S A 105, 3011–3016 (2008).

27. T. Yokosuka et al., Programmed cell death 1 forms negative costimulatory microclusters that directly inhibit T cell receptor signaling by recruiting phosphatase SHP2. J Exp Med 209, 1201–1217 (2012).

28. E. Hui et al., T cell costimulatory receptor CD28 is a primary target for PD-1-mediated inhibition. Science 355, 1428–1433 (2017).

29. B. Wang, et al., Combination cancer immunotherapy targeting PD-1 and GITR can rescue CD8(+) T cell dysfunction and maintain memory phenotype. Sci Immunol 3, (2018).

30. E. A. Philips, et al., Transmembrane domain-driven PD-1 dimers mediate T cell inhibition. Sci Immunol 9, eade6256 (2024).

31. J. L. Riley, PD-1 signaling in primary T cells. Immunol Rev 229, 114–125 (2009).

32. A. Chaudhri et al., PD-L1 Binds to B7-1 Only In Cis on the Same Cell Surface. Cancer Immunol Res 6, 921–929 (2018).

33. D. Sugiura et al., Restriction of PD-1 function by cis-PD-L1/CD80 interactions is required for optimal T cell responses. Science 364, 558–566 (2019).

34. Y. Zhao et al., PD-L1:CD80 Cis-Heterodimer Triggers the Co-stimulatory Receptor CD28 While Repressing the Inhibitory PD-1 and CTLA-4 Pathways. Immunity 51, 1059–1073.e1059 (2019).

35. S. A. Oh et al., PD-L1 expression by dendritic cells is a key regulator of T-cell immunity in cancer. Nat Cancer 1, 681–691 (2020).

36. K. Magiera-Mularz et al., Human and mouse PD-L1: similar molecular structure, but different druggability profiles. iScience 24, 101960 (2021).

37. X. Cheng et al., Structure and interactions of the human programmed cell death 1 receptor. J Biol Chem 288, 11771–11785 (2013).

38. Y. Zhao et al., PD-L1:CD80 Cis-Heterodimer Triggers the Co-stimulatory Receptor CD28 While Repressing the Inhibitory PD-1 and CTLA-4 Pathways. Immunity 51, 1059–1073 e1059 (2019).

39. Y. Zhao et al., Antigen-Presenting Cell-Intrinsic PD-1 Neutralizes PD-L1 in cis to Attenuate PD-1 Signaling in T Cells. Cell Rep 24, 379–390.e376 (2018).

40. T. Okazaki, A. Maeda, H. Nishimura, T. Kurosaki, T. Honjo, PD-1 immunoreceptor inhibits B cell receptor-mediated signaling by recruiting src homology 2-domain-containing tyrosine phosphatase 2 to phosphotyrosine. Proc Natl Acad Sci U S A 98, 13866–13871 (2001).

41. Y. N. Chen et al., Allosteric inhibition of SHP2 phosphatase inhibits cancers driven by receptor tyrosine kinases. Nature 535, 148–152 (2016).

42. E. Hui, R. D. Vale, In vitro membrane reconstitution of the T-cell receptor proximal signaling network. Nat Struct Mol Biol 21, 133–142 (2014).

43. J. Celis-Gutierrez et al., Quantitative Interactomics in Primary T Cells Provides a Rationale for Concomitant PD-1 and BTLA Coinhibitor Blockade in Cancer Immunotherapy. Cell Rep 27, 3315–3330 e3317 (2019).

44. B. Murrell et al., Gene-wide identification of episodic selection. Mol Biol Evol 32, 1365–1371 (2015).

45. J. O. Wertheim, B. Murrell, M. D. Smith, S. L. Kosakovsky Pond, K. Scheffler, RELAX: detecting relaxed selection in a phylogenetic framework. Mol Biol Evol 32, 820–832 (2015).

46. V. C. Xie, J. Pu, B. P. Metzger, J. W. Thornton, B. C. Dickinson, Contingency and chance erase necessity in the experimental evolution of ancestral proteins. Elife 10, (2021).

47. G. C. Finnigan, V. Hanson-Smith, T. H. Stevens, J. W. Thornton, Evolution of increased complexity in a molecular machine. Nature 481, 360–364 (2012).

48. N. M. Foley et al., A genomic timescale for placental mammal evolution. Science 380, eabl8189 (2023).

49. S. Alvarez-Carretero et al., A species-level timeline of mammal evolution integrating phylogenomic data. Nature 602, 263–267 (2022).

50. P. Zhou et al., Single-cell CRISPR screens in vivo map T cell fate regulomes in cancer. Nature 624, 154–163 (2023).

51. I. Siddiqui et al., Intratumoral Tcf1+PD-1+CD8+ T Cells with Stem-like Properties Promote Tumor Control in Response to Vaccination and Checkpoint Blockade Immunotherapy. Immunity 50, 195–211.e110 (2019).

52. S. J. Im et al., Defining CD8+ T cells that provide the proliferative burst after PD-1 therapy. Nature 537, 417–421 (2016).

53. R. He et al., Follicular CXCR5-expressing CD8(+) T cells curtail chronic viral infection. Nature 537, 412–428 (2016).

54. B. C. Miller et al., Subsets of exhausted CD8+ T cells differentially mediate tumor control and respond to checkpoint blockade. Nat Immunol 20, 326–336 (2019).

55. B. T. Fife, K. E. Pauken, The role of the PD-1 pathway in autoimmunity and peripheral tolerance. Ann N Y Acad Sci 1217, 45–59 (2011).

56. F. Martins et al., Adverse effects of immune-checkpoint inhibitors: epidemiology, management and surveillance. Nat Rev Clin Oncol 16, 563–580 (2019).

57. G. Myers, Immune-related adverse events of immune checkpoint inhibitors: a brief review. Curr Oncol 25, 342–347 (2018).

58. K. Adam, A. Iuga, A. S. Tocheva, A. Mor, A novel mouse model for checkpoint inhibitor-induced adverse events. PLoS One 16, e0246168 (2021).

59. H. Nishimura, M. Nose, H. Hiai, N. Minato, T. Honjo, Development of lupus-like autoimmune diseases by disruption of the PD-1 gene encoding an ITIM motif-carrying immunoreceptor. Immunity 11, 141–151 (1999).

60. H. Nishimura et al., Autoimmune dilated cardiomyopathy in PD-1 receptor-deficient mice. Science 291, 319–322 (2001).

61. T. Yokosuka et al., Spatiotemporal regulation of T cell costimulation by TCR-CD28 microclusters and protein kinase C theta translocation. Immunity 29, 589–601 (2008).

62. M. A. Mintz et al., The HVEM-BTLA Axis Restrains T Cell Help to Germinal Center B Cells and Functions as a Cell-Extrinsic Suppressor in Lymphomagenesis. Immunity 51, 310–323.e317 (2019).

63. U. Lorenz, SHP-1 and SHP-2 in T cells: two phosphatases functioning at many levels. Immunol Rev 228, 342–359 (2009).

64. M. Marasco et al., Molecular mechanism of SHP2 activation by PD-1 stimulation. Sci Adv 6, eaay4458 (2020).

## Supplementary References

65. D. Conant et al., Inference of CRISPR Edits from Sanger Trace Data. CRISPR J 5, 123–130 (2022).

66. F. A. Wolf, P. Angerer, F. J. Theis, SCANPY: large-scale single-cell gene expression data analysis. Genome Biol 19, 15 (2018).

67. I. Tirosh et al., Dissecting the multicellular ecosystem of metastatic melanoma by single-cell RNA-seq. Science 352, 189–196 (2016).

68. X. Su, J. A. Ditlev, M. K. Rosen, R. D. Vale, Reconstitution of TCR Signaling Using Supported Lipid Bilayers. Methods Mol Biol 1584, 65–76 (2017).

69. A. D. Edelstein et al., Advanced methods of microscope control using muManager software. J Biol Methods 1, (2014).

70. J. Schindelin et al., Fiji: an open-source platform for biological-image analysis. Nat Methods 9, 676-682 (2012).

71. S. Kumar et al., TimeTree 5: An Expanded Resource for Species Divergence Times. Mol Biol Evol 39, (2022).

72. R. Allio et al., OrthoMaM v12: a database of curated single-copy ortholog alignments and trees to study mammalian evolutionary genomics. Nucleic Acids Res 52, D529–D535 (2024).

73. K. Katoh, D. M. Standley, MAFFT multiple sequence alignment software version 7: improvements in performance and usability. Mol Biol Evol 30, 772–780 (2013).

74. V. Hanson-Smith, B. Kolaczkowski, J. W. Thornton, Robustness of ancestral sequence reconstruction to phylogenetic uncertainty. Mol Biol Evol 27, 1988–1999 (2010).

75. Z. Yang, PAML 4: phylogenetic analysis by maximum likelihood. Mol Biol Evol 24, 1586–1591 (2007).

